# Extensive photophysiological variation in wild barley is linked to environmental origin

**DOI:** 10.1101/2025.03.28.645825

**Authors:** Matthieu Breil-Aubert, Katie Shaw, Jessica Royles, Cris Sales, Julia Walter, Georgia Taylor, Richard Vath, Eyal Bdolach, Lalit Dev Tiwari, Jyotirmaya Mathan, Tracy Lawson, Eyal Fridman, Johannes Kromdijk, John N Ferguson

## Abstract

- Intraspecific variation between crop wild relatives (CWRs) represents a source of untapped genetic diversity for crop improvement. At the same time, improving photosynthesis in crops has the potential to enhance yield. Thus, exploring variation for photophysiology within CWRs is an important, yet underexplored, research area.
- We describe a common garden experiment where 320 wild barley accessions were grown across two seasons. A photophysiology phenotyping pipeline was employed to quantify >30 traits within this diversity panel. Population genetics, genome-wide association analyses (GWAS) and deep phenotyping were performed to address local adaptation hypotheses.
- Heritable variation was detected across this photophysiological spectrum, with genotype-by-environment (GxE) interactions being prevalent. Evidence for local adaptation was observed in the form of subpopulation differences, signals of selection, and allele frequency variation associated to markers identified via GWAS. Phenotyping of representative accessions across distinct water availabilities highlighted a role for stomatal conductance (*g*_s_) in adaptation to dry environments.
- We identified substantial variation in key photosynthesis-associated traits in a CWR closely related to barley, an economically important crop species. Our results demonstrate that this variation is partially due to local adaptation, where plasticity in *g*_s_ appears important for maintaining photosynthesis and biomass accumulation in water restricted conditions.

## Introduction

Barley (*Hordeum vulgare*) is a globally important crop species especially within Europe where it is one of the most productive crops in the United Kingdom and the European Union. Currently, barley is predominately grown for animal feed and malting; however, given its potential as a more nutritious alternative for cereal-based food production that wheat, rice or maize cultivation for human consumption is increasing (Baik & Ullrich, 2008; Meints *et al*., 2016). Depending on the agronomic environment, genetic gain for yield in barley ranges anywhere from 0.43-to-1.07% per annum (Cossani *et al*., 2022; Giménez *et al*., 2024; Åstrand *et al*., 2024). These rates of yield increases will be insufficient to meet future demands, especially considering both the renewed interest in barley for food production and the challenges posed by climate change (Åstrand *et al*., 2024). Consequently, breeding barley varieties for improved yields that are stable in the face of reduced resource inputs and environmental stress is a key priority for future food security (Jiang *et al*., 2024).

The efficiency with which radiative energy intercepted by crop canopies is converted into biomass by photosynthesis is rarely close to the theoretical maximum (Zhu *et al*., 2010; Slattery & Ort, 2015). Furthermore, across the major non-leguminous C_3_ species this conversion efficiency is lowest on average in barley (Slattery & Ort, 2015). Photosynthetic conversion efficiency is a major determinant of yield potential (Monteith *et al*., 1997), thus improving photosynthesis has become increasingly recognised as viable target for increasing crop yields (Long *et al*., 2006; Zhu *et al*., 2010). Here, there are several strategies available given the numerous limitations that may limit photosynthesis under agronomic conditions (Reviewed by: Croce *et al*., 2024). One approach that is receiving increasing consideration is to explore natural variation in photosynthesis as a source of novel genes and alleles that could help finetune photosynthesis to specific environmental conditions and eventually enhance yield (Reviewed by: Theeuwen *et al*., 2022). To this end, it is pertinent here to consider the recent work of Gao *et al*., (2024) who evaluated multiple photosynthesis-related parameters via chlorophyll fluorescence in the field across 23 spring barley varieties over distinct developmental stages. Here, traits that define the rate at which atmospheric CO_2_ can be assimilated, specifically the maximum efficiency of photosystem II in the light (ΦPSII) and non-photochemical quenching (NPQ), demonstrated moderate-to-high heritabilities depending on the developmental stage being phenotyped. The presence of a discernible genetic basis to these traits highlights their amenability for breeding. Moreover, Gao *et al*., (2024) observed that these two parameters were significantly correlated with yield-related parameters, providing empirical evidence to support the theory that improving photosynthesis in barley is a feasible means to improve yield.

Like all of our most important crop species, barley has largely been bred and cultivated in benign, agronomically managed environments (Jiang *et al*., 2024). This is in stark contrast to crop wild relatives (CWRs), i.e., wild plant species that are related to crops, but that have not passed through a domestication bottleneck and that persist across a range of environmentally challenging habitats (Tanksley & McCouch, 1997). As a result, these CWRs maintain much higher degrees of genetic variability between distinct ecotypes compared to their domesticated relatives and have been challenged in much harsher environments over thousands of years (Zhang *et al*., 2017). Consequently, CWRs represent an untapped resource of genetic variation for improving photosynthesis, including under environmentally challenging situations. Several recent studies provide evidence in support of this approach. For example, McAusland *et al*., (2020) demonstrated that many wild wheat relatives are capable of maximal rates of photosynthesis that are significantly higher than those achievable by modern common wheat varieties. Similar findings were also made by a studying focusing on barley landraces (Stevens *et al*., 2021). Also, a CWR from the Brassicaceae family, *Hirschfelida incana*, has recently been identified to have exceptionally high photosynthetic light-use efficiency that far exceeds that of closely related *Brassica* crop species (Garassino *et al*., 2022).

With specific reference to barley, there are now excellent opportunities to begin to leverage the variation that exists in CWRs to improve photosynthesis in elite barley varieties. For example, the recently published pangenome of barley incorporates 23 wild barley genomes alongside 53 domesticated barley genomes (Jayakodi *et al*., 2024) which allows easy identification of novel structural variants of interest that have been lost due to domestication and could accelerate their re-integration into domesticated barley from wild barley. Additionally, there are now two established and well sequenced diversity collections of wild barley that permit genome-wide association studies (GWAS) to identify marker-trait associations (Prusty *et al*., 2021; Sallam *et al*., 2024). These diversity sets have already been used to identify novel genes underlying disease resistance, phenology, lemma color and circadian clock rhythmicity (Prusty *et al*., 2021; Sallam *et al*., 2024).

The Barley 1K (B1K) collection of wild barley (first described in: Hübner *et al*., 2009) is an ideal model system for studying local adaptation. This diversity set comprises over 1000 distinct accessions collected from a relatively small geographic range across the Southern Levant. Despite this, distinct genetic clusters exist among these accessions, where there is minimal gene flow due to geographic barriers. Environmental variables (e.g., soil water capacity, temperature and elevation) have been shown to explain a significant proportion of the total genetic and phenotypic variation across subsets of the B1K collection (Hübner *et al*., 2009; Chang *et al*., 2022). Additionally, the B1K collection has been used to identify important loci that underly the response of circadian clock rhythmicity to heat stress (Prusty *et al*., 2021; Tiwari *et al*., 2024), where associated allelic variation is also linked to fitness plasticity in response to heat stress (Tiwari *et al*., 2024). Collectively, these existing studies serve as an encouraging precursor for studying local adaptation and natural variation of photosynthesis within wild barely as a CWR.

Local adaptation is widely recognised phenomenon (Reviewed by: Hereford, 2009), but the traits and genes involved are often unknown. To this end, there have been very few studies that have explored the role photosynthesis and associated processes play in local adaptation. Moreover, these studies tend to be concentrated on the model species Arabidopsis (*Arabidopsis thaliana*). For example, Elfarargi *et al*., (2023) demonstrated that a population of Arabidopsis that has colonised an island characterised by fog-based precipitation had much higher rates of stomatal conductance (*g*_s_) compared to its most closely related outgroup from a temperate region of Africa. Here, an allele promoting increased stomatal opening was observed to be under positive selection on the colonised island to promote local adaptation to humid conditions via an anisohydric strategy. Taking a different approach, Oakley *et al*., (2018) employed a recombinant inbred line (RIL) population derived from Swedish and Italian ecotypes to determine that quantitative trait loci (QTL) regulating the photosynthetic response to cold stress co-localised with QTL regulating local adaptation based on reproductive fitness.

The study presented in this paper extends this approach by describing the most comprehensive screening of photosynthesis across natural accessions of a CWR yet performed. For this we utilised 320 accessions from the B1K diversity collection. We reveal extensive, heritable variation across traits defining light-saturated photosynthesis, the response of photoprotection and PSII quantum yields to dynamic irradiance, and limitations to light-saturated photosynthesis. We provide evidence to suggest that this variation is partially a result of differential selection across the various sub-populations for differing photo-physiological properties. Specifically, we highlight how *g*_s_ may play a key role in facilitating local adaptation to drier environments. More broadly, this research showcases that extensive variation in photosynthesis is present in wild barley and that some of this variation is comparatively greater than what has recently been demonstrated in domesticated barley (Gao *et al*., 2024).

## Materials and Methods

### Plant material and common garden experiments

This study incorporated 320 accessions from the Barley1K (B1K) collection of wild barley (*Hordeum spontaneum*) collected from 51 sites across Israel (Elfarargi *et al*., 2023). Common garden experiments incorporating these accessions were carried out in 2021 and 2022 at the National Institute of Agricultural Botany (NIAB, Cambridge, UK). The sites of the two common garden experiments were ∼870 m apart across the two years (Supporting Figure S1). In 2021, all 320 accessions were included in the common garden experiment (Supporting Table S1). In 2022, we included a subset of 270 accessions, where all 270 were present in the 2021 experiment (Supporting Table S1). In both years, an alpha lattice twice replicated design was employed for the common garden experiments, such that each accession was represented by two plots. Each replicate of the experiment consisted of eight blocks of 40 plots in 2021 and six blocks of 45 plots in 2022. Each plot was separated by 30 cm from adjacent plots. In both years, plots consisted of four rows. The outer two rows were made up of drilled spring barley (cv. Laureate) with the two inner rows being hand-transplanted wild barley. Rows were spaced 10 cm apart and were 40 cm long. Each inner row of wild barley consisted of eight transplanted plants. Before transplanting, the wild barley was hand sown in modular trays of M2 potting compost for germination in an ambient temperature glasshouse. Following germination, the wild barley was then moved into a vernalisation room set to 5 °C for approximately one month. Before transplanting, the plants were moved outside into a hardstanding area to acclimate to outside conditions. In both years, transplanting into the common garden occurred in late April. Precise dates of drilling, sowing, vernalisation and transplanting are provided in Supporting Table S2 alongside agronomic inputs.

### Phenotyping pipeline

We developed a high-throughput phenotyping pipeline to screen a multitude of traits associated with photosynthetic performance. This pipeline is demonstrated in a flow chart shown in Supporting Figure S2 and described below.

All phenotyping was scheduled to co-occur as close as possible to heading date. Heading date was scored daily on a plot-by-plot basis as the emergence of the spike out of the flag leaf sheath (Zadoks *et al*., 1974). A plot was considered to be heading if more than 50% of the wild barley plants within that plot were heading. Days to heading (DTH) was then calculated as the number of days between the date of transplanting of the seedlings into the common garden experiment and the date of heading.

Three plants were phenotyped per plot, meaning six plants were phenotyped in total for each accession within a year. Plants to be phenotyped were flagged with a barcoded tag. On the day before phenotyping, flagged plants were cut at the base of the main stem with the cut end immediately placed into water. Excised stems were then returned to the laboratory and recut under water into individual 10 ml centrifuge tubes to maintain the water column. Stem harvesting was completed between 1500 and 1700 after which the excised stems were left on the laboratory bench at room temperature overnight. Phenotyping proceeded the next day at 0630 and continued until 1430. We have previously shown that all phenotyping performed on these excised stems in barley generates data that is comparable to phenotyping leaves attached to the whole plant (Ferguson *et al*., 2023b).

Light-saturated gas exchange of the penultimate leaf was measured using LI-6400XT infra-red gas analysers (LI-COR Inc., Lincoln, NE) equipped with 6400-40 leaf chamber fluorometer LED light sources. Before measuring gas exchange, leaves were light acclimated under a series of LED panels for 30 minutes. The LED panels were set to a saturating irradiance of 1800 µmol m^-2^ s^−1^ photosynthetically active radiation (PAR) at leaf level to match the PAR setpoint in the leaf chambers of the infra-red gas analysers. The remaining conditions in the leaf chambers were set as follows: 25°C block temperature, 400 µmol s^−1^ air flow, 65-75% relative humidity (RH), and 400 µmol mol^-1^ reference CO_2_ concentration. Once moved from under the LED panels and into the leaf chambers, gas exchange was logged every 10 seconds for 15 minutes. The mean values from the last two minutes were taken for light saturated photosynthesis (*A*_sat_) and stomatal conductance to water vapour (*g*_s_), which were also used to calculate intrinsic water use efficiency (iWUE) as *A*_sat_/*g*_s_. In 2022, we additionally performed a *mini* light response (*A*-Q) curve following this measurement of light saturated gas exchange. Here, a program was immediately initiated after the 15 minutes of high light to incrementally drop the light intensity in the following steps: 1800, 1100, 500, 300, 150, and 50. Two minutes after each light step, photosynthesis was logged before dropping to the next light level.

A randomly selected subset of accessions (Supporting Table S2) was selected for phenotyping the response of photosynthesis to changes in the intracellular concentration of CO_2_ (*A*-*C*_i_ curve). All accessions used for *A*-*C*_i_ curves in 2021 were also used in 2022 with some additional accessions added in 2022. These measurements were performed using LI-6800 infra-red gas analysers equipped with a standard 6cm^2^ leaf chamber (LI-COR Inc., Lincoln, NE) with environmental conditions set as follows: 25°C temperature, 400 µmol s^−1^ air flow, 65% relative humidity (RH), 400 µmol mol^-1^ reference CO_2_ concentration and 1800 µmol m^-2^ s^−1^ PAR. Once stomatal conductance and photosynthesis were stable, an *A*-*C*_i_ curve program was initiated to log rates of gas exchange at the following CO_2_ reference concentrations: 400, 300, 200, 100, 50, 400, 400, 700, 1000, 1300, and 1800 µmol mol^−1^ waiting between 1.5-3 minutes between each CO_2_ step depending on standard stability criteria. Where leaves did not fill the cuvette completely, the leaf area contained in the leaf cuvette was calculated from leaf width measurements and adjusted accordingly in the gas exchange calculations.

The above-described gas exchange measurements were performed on the penultimate leaf (i.e. the leaf below the flag leaf). Following gas exchange phenotyping, the penultimate leaf was excised and flattened under a glass sheet. The leaf was then photographed using a mounted digital camera. The leaf was carefully folded into a coin envelope and placed into a drying oven set at 60°C for seven days. The photographs of the penultimate leaves were used to measure leaf area using the Easy Leaf Area software (Easlon & Bloom, 2014). Once the same leaves were fully dried, we then calculated specific leaf area as the ratio of the leaf area to dry mass.

Finally, an approximately 3 x 1 cm strip of tissue from the flag leaf was cut and used for chlorophyll fluorescence in a FluorCam closed chlorophyll fluorescence imaging system (Photon Systems Instruments, Brno, Czechia) exactly as described previously (Ferguson *et al*., 2023a). Briefly, we placed leaf samples on top of damp filter paper encased between non-reflective glass plates and dark adapted overnight by covering in aluminium foil. We then measured the quantum efficiency of PSII (*F*_v_/*F*_m_) as well as the response of non-photochemical quenching (NPQ) and photosystem II (PSII) operating efficiency (□PSII) to an actinic light (1800 µmol m^-2^ s^−1^) being switched on for 600 seconds and then off 800 seconds.

An approximately 3 x 1.5cm strip of tissue was excised from the dried leaves used to calculate SLA and ground in a bead mill. The ground, dried tissue was precisely weighed (0.5mg +/- 10%) into individual tin capsules and underwent analysis for leaf carbon (%C) and leaf nitrogen (%N) as a percentage of dried mass as well as carbon isotope composition (δ^13^C) and nitrogen isotope composition (δ^13^N). These measurements were carried out using a Costech Elemental Analyser attached to a Thermo DELTA V mass spectrometer via Conflo IV in continuous flow mode at The Godwin Laboratory for Palaeoclimate Research (Cambridge University, UK).

### Photosynthesis modelling

The light-response (*A*_N_-*Q*) data generated in 2022 was fitted using the custom fit_AQ_curve() function available on the “AQ_curves” github repository (Tomeo, 2024). This function uses the non-rectangular hyperbola model described in Lobo *et al* (2013). For this model, we set the convexity parameter which regulates the curvature of the response of photosynthesis to light at 0.7 following von Caemmerer (2000). From this, we obtained estimates for respiration in the light (R_L_) and the apparent maximum quantum yield (ΦCO_2max_).

The CO2-response (*A*-*C*_i_) data were fitted according to the FvCB model (Farquhar *et al*., 1980) using the function fitacis() function from the plantecophys R package (Duursma, 2015). We used the bilinear method to estimate transition points. From this, estimates of the maximum rate of Rubisco carboxylation (Vcmax), the maximum rate of electron transport for RuBP regeneration (Jmax) and triose phosphate utilisation (TPU) limitation (TPU) were obtained on *c*_i_ basis. We additionally utilised the *A*-*C*_i_ data to estimate the stomatal limitation (SL) on photosynthesis following (Long & Bernacchi, 2003).

We used linear and exponential models to describe the induction of NPQ in response to the actinic light being switched on. We also used an exponential model to describe the relaxation of NPQ and recovery of □PSII in response to the actinic light being switched off. We have described these models previously (Ferguson *et al*., 2023a). These models allowed us to determine the following: the slope of the initial induction of NPQ (NPQ_linear_); the amplitude (NPQ_ind-amp_) and rate (NPQ_ind-rate_) of NPQ induction; the amplitude (NPQ_rel-amp_), rate constant (NPQ_rel-rate_) and model offset (NPQ_rel-res_) of NPQ relaxation; the amplitude (□PSII_rec-amp_), rate (□PSII_rec-rate_), and model offset (□PSII_rec-res_)of □PSII recovery.

### Statistical analyses

Unless stated all data handling and statistical analyses were performed within R (R Core Team, 2021) using ggplot2 (Wickham, 2009) for graphing.

We used the H2cal() function from the inti R package (Lozano-Isla *et al*., 2024) to generate breeding values (Best Linear Unbiased Predictors (BLUPs) and Best Linear Unbiased Estimator (BLUEs). Mixed models were constructed that incorporated genotype, block, column, replicate and days post heading as fixed (BLUPs) or random (BLUEs) predictors of trait values on a year-by-year basis. We also produced joint-year models that include interactions between these predictors and year. The variance components from these models were used to calculate broad sense heritability (*H*^2^) according to Cullis *et al* (2006) for BLUPs (*H*^2^_C_) and Piepho & Möhring (2007) for BLUEs (*H*^2^_PM_).

Pairwise trait correlations and correlations between the same traits across years were examined via the Pearson correlation coefficient (*r*) and associated p-values (*p*) using the cor() base R function. Significant correlations were defined as those where *p* ≤ 0.05. Differences in traits between subpopulations were tested via one-way analysis of variance (ANOVA) comparison of means testing using the aov() based R function. Significance differences between subpopulations were defined as those where *p* ≤ 0.05. Post-hoc Tukey tests were performed to determine which subpopulations were significantly different from one another. This was achieved using the HSD.test() function from the agricolae R package (Mendiburu & Simon, 2015).

A quantitative estimate of phenotypic plasticity (GxE) was estimated by calculating the difference in trait values for each accession between field years, weighted by the average population value for each respective year. This was computed using the formula: (Population Mean 2021 + BLUE accession value 2021) - (Population Mean 2022 + BLUE accession value 2022). This approach captures the extent of phenotypic variation in response to environmental differences between years.

### Population genetics

A previously published customised SNP genotyping dataset for wild barley (Tiwari *et al*. 2024b) was used for population genetics and genome-wide association analyses. accessions from this recent study are common with our study (Supporting Table S1). We utilised the SNP data to perform population genetics analyses specific to the accessions included in our study as described later.

For phylogenetic analyses, we first converted the VCF format of the previously described SNP dataset to a ‘genind’ object using the loci2genind function from the pegas package (Paradis, 2010). We then calculated a neighbor-joining tree (dendrogram) with bootstrap support based on Nei’s distance, using the aboot function from the poppr package (Kamvar *et al*., 2015). Finally, we extracted the genotype order from the tree to properly align and sort the structure plot.

For STRUCTURE analyses, we first converted the VCF format to a numeric matrix using a custom function. To estimate the optimal number of subpopulations (dimensions), we used the estimate_d function from the alstructure package (Cabreros & Storey, 2019), which is based on the method from (Leek, 2011). We then applied the alstructure function from the same package to compute global ancestry estimates under the admixture model, utilizing the ALStructure algorithm (Cabreros & Storey, 2019). A genotype was assigned to a subpopulation if it had an admixture proportion greater than 50% in one “cluster” (subpopulation).

We performed analyses to detect signals of natural selection using the DRIFTSEL R package (Karhunen *et al*., 2013). DRIFTSEL is a Bayesian method that compares predicted and observed mean additive genetic values to generate the S statistic, which indicates whether population divergence is driven by divergent selection (S ∼ 1), stabilizing selection (S ∼ 0), or genetic drift (intermediate S values). This method is particularly effective for small datasets and can differentiate between drift and selection even when *Q*_ST_ (divergence in quantitative traits) *F*_ST_ (divergence in neutral molecular markers) are equal, assuming that phenotypic variation is determined by additive genotypic variation. To estimate the coancestry coefficient matrix, we used the RAFM package (Karhunen & Ovaskainen, 2012). Both the RAFM and DRIFTSEL models were fit using 15,000 Markov chain Monte Carlo (MCMC) iterations, with the first 5,000 iterations discarded as burn-in and the remaining samples thinned by a factor of 2, resulting in 5,000 posterior distribution samples. Due to a lack of information on the dams and sires of the phenotyped wild barley individuals, we made a slight modification to RAFM and DRIFTSEL. We adopted a conservative assumption that each genotype’s dam and sire are the same and that all genotypes are unrelated.

We carried out a genome-wide association study (GWAS) using three independent, iterative statistical models to enhance accuracy. These models included the multi-locus mixed-model (MLMM; (Segura *et al*., 2012)), the Bayesian-information and Linkage-disequilibrium Iteratively Nested Keyway (BLINK; (Huang *et al*., 2019)), and the Fixed and Random Model Circulating Probability Unification (FarmCPU; (Liu *et al*., 2016b)). All analyses were performed in R using GAPIT Version 3 (Wang & Zhang, 2021) applied to both BLUEs and BLUPs. These GWAS models are described in the associated references, but all included cofactors to account for population structure and kinship. To prioritise “high-confidence” marker trait associations, herein termed quantitative trait loci (QTL), we retained only those identified with both BLUEs and BLUPs across a joint-years model for further analysis. The genome-wide statistical significance threshold for all methods was set using the Bonferroni correction (α = 0.05), which adjusts the threshold according to the number of SNPs tested.

Alongside comparing mean trait values (see above), we also compared allele frequencies of SNPs that passed the above-described significance threshold across subpopulation. This was achieved by converting the genotype into allele frequencies with homozygous references alleles coded as 0 and homozygous alternate alleles coded as 1.

### Targeted experiment on representative genotypes from Steppe Jerusalem and Desert Jordan subpopulations

To test hypotheses regarding the adaptive capacity to reduced water availability, we selected six accessions for phenotyping under distinct water availabilities. These accessions were selected based on principal component analyses (PCA) performed using prcomp(). The PCA biplot was visualised using fviz_eig() (Kassambara & Mundt, 2020). The traits used for the PCA were leaf area, SLA, *A*_sat_, *g*_s_, δ^13^C, δ^15^N, DTH, final NPQ, NPQ_rel-amp_, and maximum NPQ. We selected six accession that well-represented the total phenotypic trait spaces (Supporting Figure S3). Three of these accessions were assigned by our STRUCTURE analysis (Figure 1B) to the “Desert Jordan” subpopulation (B1K-05-12, B1K-05-08, B1K-12-10) and three were assigned to the “Steppe Jerusalem” subpopulation (B1K-17-17, B1K-10-01, B1K-49-10).

**Figure 1.**
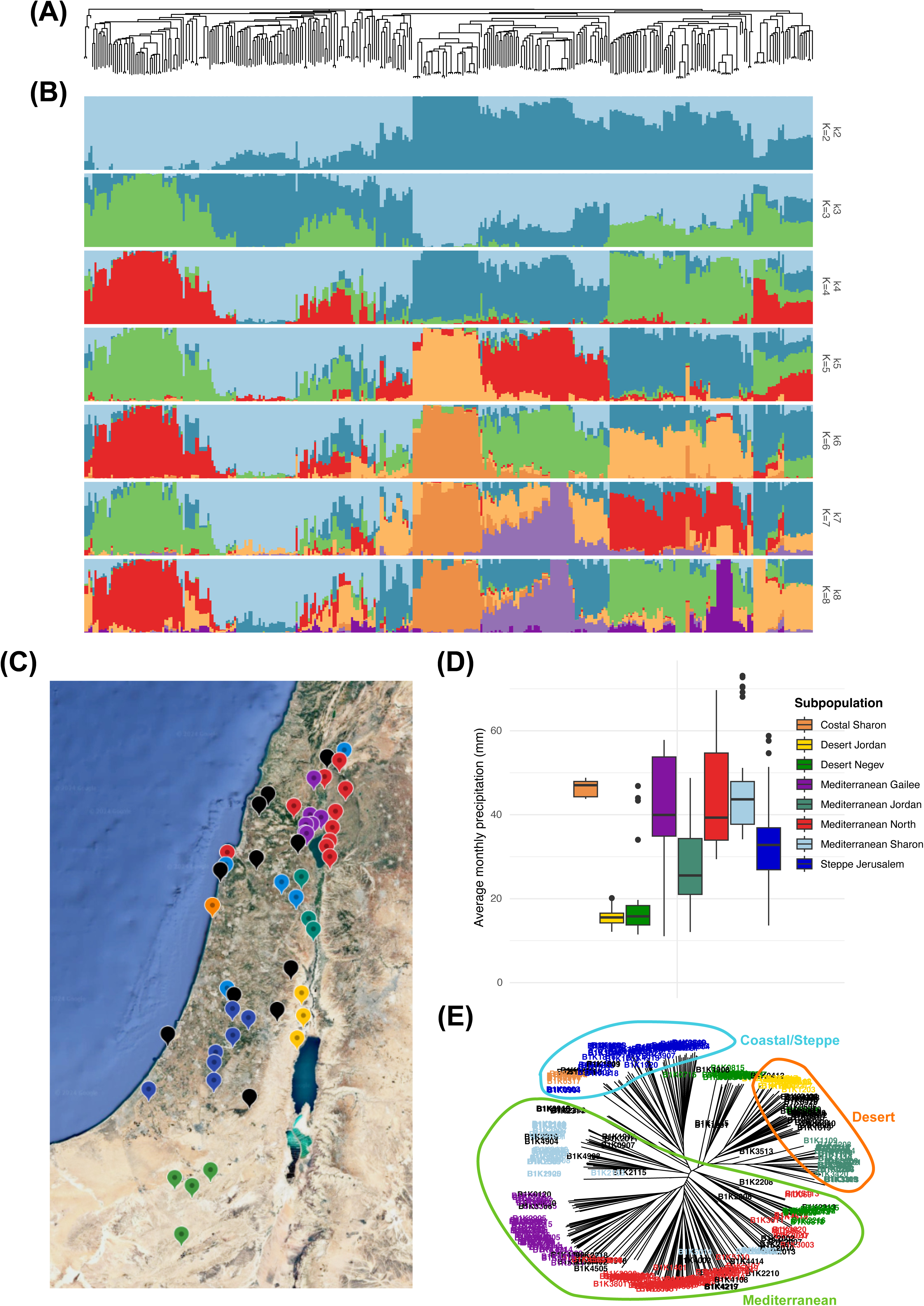
Population genetic structure of 320 wild barley accessions across Israel. (A) Unrooted neighbor-joining (NJ) tree computed by Nei distance (B) Results of the STRUCTURE analyses at *K =* 2-8. (C) Map showing the geographic sites where accessions were collected. Sites are coloured according to the STRUCUTRE-defined subpopulations that the associated accessions belong to. (D) Difference of year precipitation between subpopulations. (E) Unrooted Neighbour-joining tree computed by Nei distance. Accessions are coloured according to the STRUCTURE-defined subpopulations they belong to. Accessions are clustered into three major groups which corresponded to broad geographic areas (Mediterranean, Desert, and Coastal/Steppe) as indicated by coloured ellipses.

Seeds from these accessions were placed in petri dishes containing damp filter paper. These were then wrapped in aluminium foil and transferred to the fridge for six days before moving to room temperature for four days. Germinated seedlings were transplanted into 1.5L pots containing 950g of topsoil. Once the plants were established, a known mass of polypropylene beads was placed on top of the soil to limit evaporation from the soil. Plants were grown under glasshouse conditions (Cambridge University Botanic Garden, Cambridge, UK). A nematode biological control treatment (*Steinernema fletiae* and *Steinernema carpocapsae,* Koppert Biological Systems, Haverhill, UK) was applied to each pot on seven days post transplanting to control scarid fly. Environmental conditions in the glasshouse were set as follows: 60% relative humidity (RH), 28°C day temperature. 23°C night temperature. Daylength was set to 16 hours. A TinyTag Ultra 2 TGU-4500 (Gemini Data Loggers, Chichester, UK) was placed in the glasshouse for the duration of the experiment to record temperature and RH (Supporting Figure S4). A BF5 sunshine sensor and GP1 data logger (DELTA-T Devices, Cambridge, UK) were also placed in the glasshouse to measure incident radiation (Supporting Figure S5).

Pots were initially well-watered for 14 days to allow for plant establishment, before being exposed to two contrasting water availability treatments. Here, the plants were either maintained at 40% relative soil water content (rSWC) or 80% rSWC, where 100% rSWC is equivalent to field capacity. Pots were maintained at target rSWCs on a daily basis as described previously (Ferguson *et al*., 2019). 18 soil-only control pots were used to account for direct evaporation from the soil and these were distributed equally among the treatment pots.

*A*-*C*_i_ response measurements were performed 34-40 days post transplanting on recently fully expanded leaves using LI-6400XT infra-red gas analysers (LI-COR, Lincoln, NE) as described previously (Ferguson *et al*., 2023b). Five plants per accession per treatment were measured between 0700 and 1600. Data were processed as described in the above “Photosynthesis modelling” section.

At 55-days post transplanting, all leaves from each plant were excised and a top-down photograph was taken to measure total leaf area using Easy Leaf Area (Easlon & Bloom, 2014).

Two-way ANOVAs were carried out to analyse the data from our glasshouse experiment. This was performed by using the aov() function in R, with genotype nested within subpopulation, to test for statistically significant differences between rWSC treatment and subpopulation (and potential interactions). The TukeyHSD() function in R was used to determine significant pairwise interactions. shapiro.test() and leveneTest() were used to test for the normality and equal variance assumptions of ANOVA. Parameters that did not meet the normality assumptions (total leaf area and iWUE) were log-transformed prior to performing further statistical analysis. Note that for total leaf area, the equal variance assumption was not met. Percentage change was determined by calculating the change in parameter mean between the 80% and 40% rWSC treatments for each sub-population, dividing by the 80% rWSC parameter mean and multiplying by 100.

## Results

### Wild barley diversity is structured in genetic clusters across Southern Levant that demonstrate distinct bioclimatic profiles

The population structure of the wild barley accessions incorporated as part of this study was investigated using STRUCUTRE. These analyses allowed us to group all accessions into subpopulations according to patterns of genetic variation (Figure 1). Through this approach we observed clear clustering of individual accessions into distinct subpopulations (Figure 1A). Using a conditional factor model, we determined that the optimum number of subpopulations (K) was eight (Figure 1B), with 69 accessions being classified as admixed (Supporting Table S1), i.e., accessions whose alleles represent mixed ancestry between subpopulations. In general, higher K values, i.e. those greater than 5, tend to be indicative of fine-scale differentiation between subpopulations (Evanno *et al*., 2005), as such our analyses suggest a high degree of subpopulation specificity. This is plausibly driven by local adaptation and reduced gene flow between genetic clusters.

In general, the subpopulations we identified tended to cluster geographically. Consequently, we named these subpopulations according to the regions where they were collected (Figure 1C, Supporting Table S1): Mediterranean North (71 accessions), Mediterranean Sharon (37 accessions), Mediterranean Galilee (41 accessions), Costal Sharon (8 accessions), Mediterranean Jordan (29 accessions), Steppe Jerusalem (48 accessions), Desert Jordan (19 accessions), and Desert Negev (33 accessions). The Mediterranean subpopulations dominate the northern and western parts of the sampling area. The Coastal and Steppe subpopulations are primarily located in transitional zones between the Mediterranean and Desert regions, while the Desert subpopulations are concentrated in the Southern and Eastern arid regions. Admixed individuals are predominantly found at the interfaces between these regions, suggesting genetic exchange in areas of overlapping environmental conditions.

To gauge the potential for climatic drivers of differentiation between the geographically distinct subpopulations, we obtained bioclimatic variables using the latitudinal and longitudinal coordinates associated with the point of collection of all accessions. We then compared these bioclimatic variables across the subpopulations (Figure 1D; Supporting Figure S6). Accessions aligned to the Desert subpopulations come from areas characterised by low precipitation and high temperatures characteristic of arid climates. Contrastingly, the point of origin of the Mediterranean accessions tended to be much wetter and milder in temperature. The regions harbouring the Coastal and Steppe accessions occupied intermediary climatic regions.

We additionally used a neighbour-joining method (Nei’s distance) to compute genetic distances and generate an unrooted phylogenetic tree of all accessions (Figure 1E). This approach corroborated our STRUCTURE analyses and confirmed the geographic clustering of the identified subpopulations. Here, the Mediterranean subpopulations form a distinct clade that is separate from the clades that incorporate the Desert and Coastal-Steppe subpopulations. This further supports the hypothesis that genetic differentiation is driven by both geographic isolation and adaptation to specific environmental conditions. Our study incorporates many unique accessions compared to the recent work of Chang et al. (2022; Supporting Table S1), however our classification of common accessions to distinct subpopulations largely aligns with their previous analyses.

### Wild barley accessions demonstrate heritable variation for photosynthetic and life history traits

We monitored daily temperature and water inputs (precipitation and irrigation) during the 2021 and 2022 common garden experiment. The 2022 growing season was markedly warmer than the 2021 growing season (Supporting Figure S7). The average daily maximum temperature in 2021 was 16.23°C (SE = 0.56°C), whereas in 2022 it was 18.98°C (SE = 0.45°C). In 2021, the average daily minimum temperature was 7.37°C (SE = 0.52°C), whereas in 2022 it was 9.18°C (SE = 0.33°C). Water inputs were similar between the two years, where the average daily combined precipitation and irrigation was 0.46mm (SE = 0.13mm) in 2021 and 0.59mm (SE = 0.18mm) in 2022 (Supporting Figure S8).

Consistent with the warmer temperatures, we saw an overall shift towards greater specific leaf area (SLA) in 2022 compared to 2021(Figure 2A). SLA can be used as a proxy for relative growth rate (Liu *et al*., 2016a), where the two are typically positively correlated and respond in the same manner to warmer temperatures that are not stressful (Loveys *et al*., 2002). Contrary to this, however, we observed a marked reduction in light-saturated photosynthetic assimilation (*A*_sat_), which was reduced on average in 2022 compared to 2021. This was matched to trend for stomatal conductance (*g*_s_), which was also reduced in 2022. The overall reduction in *A*_sat_ and *g*_s_ across all accessions was similar in magnitude, which is reflected in intrinsic water use efficiency (*iWUE*) variation being similar in 2021 and 2022.

**Figure 2.**
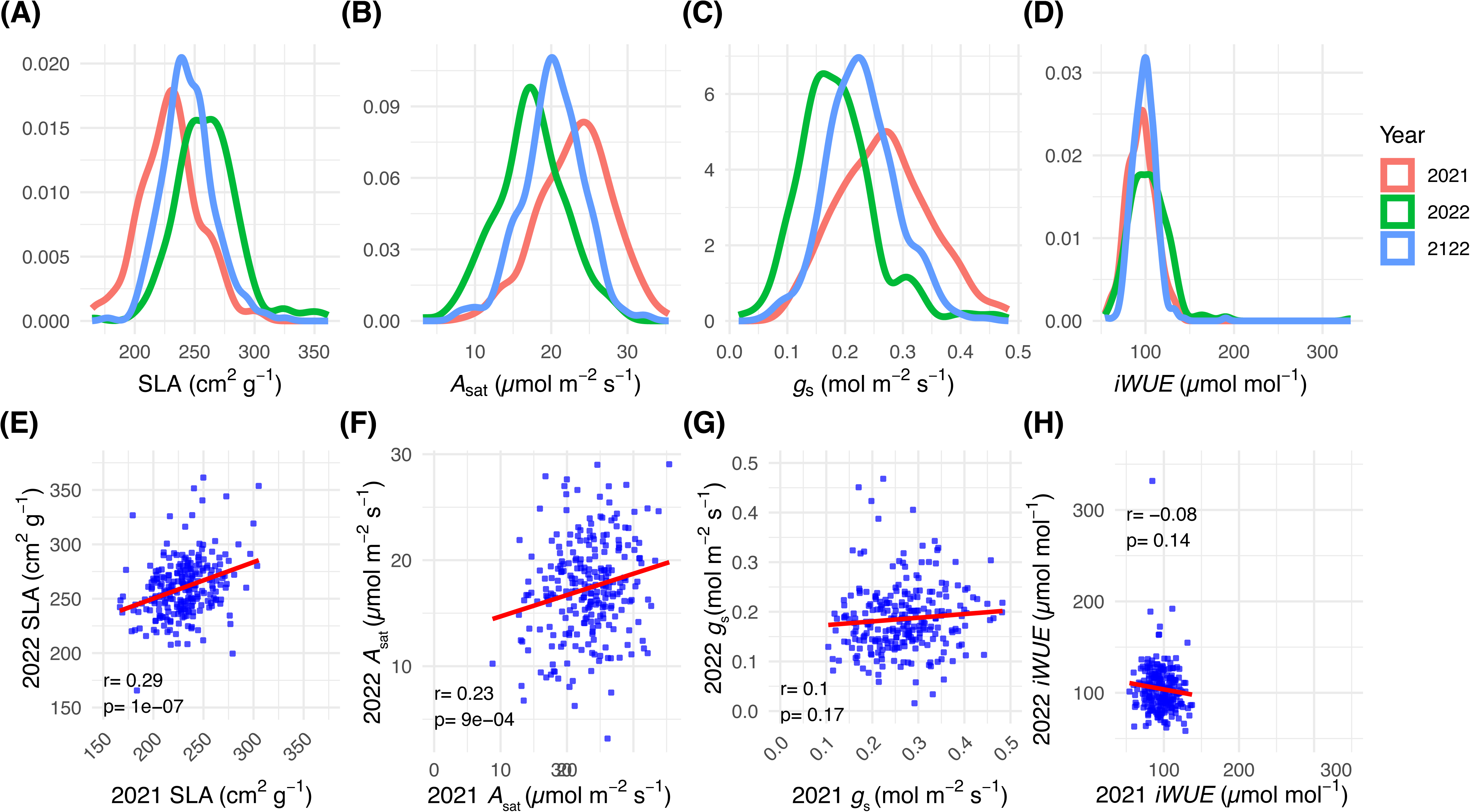
Phenotypic BLUEs variation characterised across the B1K diversity set. (A-D) Histograms showing variation for specific leaf area, photosynthesis, stomatal conductance and intrinsic water use efficiency. Histograms are coloured according the models the BLUEs are derived from, i.e. 2021 model, 2022 model, or joint year model (2121). (E-H) Correlations between specific leaf area, photosynthesis, stomatal conductance and intrinsic water use efficiency between 2021 and 2022 field trials.

The shift in variation for the above-described traits across each of the common garden experiments was not consistent on an accession-by-accession basis, since variation for most traits was only weakly correlated across the two experiments. SLA was an exception to this general trend, where accessions that demonstrated high SLA in 2021 tended to also do so in 2022 and vice versa (Figure 2E). The same was also true of heading date (Supporting Figure S9), although we did note that a subset of accessions demonstrated much earlier heading in 2022 (Supporting Figure S10), which caused the distribution to appear different when plotted (Supporting Figure S9). In general, these results suggest that variation in growth rate and phenology was strongly influenced by underlying genetics in these two environments. This suggestion is supported by the extent of observed variation explained by the accession term in the mixed linear models used to partition variances for these traits (Supporting Table S3). Variation in *A*_sat_ was marginally correlated between the two common garden experiments (Figure 2F). However, *g_s_* and *iWUE* were not correlated at all (Figures 2G-H), which suggests there were strong genotype-by-environment (GxE) interactions for water-use in these accessions (Figures 2C-D; Supporting Figure S10). Traits related to the light-dependent photosynthetic processes demonstrated moderate correlations between the two experiments (Supporting Figure S10).

We calculated heritability in order to estimate the extent to which the overall variation could be associated to genetic variation. Average heritability estimated according to Piepho & Möhring (2007; *H^2^*_PM_)) using the joint-year data was 0.37 for all traits (Table 1). The area of the penultimate leaf showed the highest *H^2^*_PM_ (0.81). The number of DTH was less heritable (0.28). Traits associated to the light-dependent photosynthetic reactions measured via chlorophyll fluorescence tended to be moderately heritable (0.29-0.59), whereas leaf gas exchange-associated traits were generally less heritable (0.01-0.49). Apart from δ^13^C (0.41), traits associated to leaf elemental composition demonstrated low heritability (0.03-0.13). Depending on the trait, *H^2^*_PM_ estimated by the individual year models was quite distinct (Supporting Table S4). Gas exchange-associated traits demonstrated higher heritability in general in 2021 compared to 2022. For example, *V*_cmax_ and *J*_max_ had *H^2^*_PM_ of 0.5 and 0.48 in 2021 and 0.21 and 0.39 in 2022. Conversely, traits estimated via chlorophyll fluorescence appeared more heritable in 2022, where, for example, maximum NPQ and final NPQ had *H^2^* values of 0.61 and 0.57 in 2021 and 0.49 and 0.38 in 2021. These findings highlight that the extent of variation imparted by the environment was different between the two growing seasons in line with substantial GxE.

**Table 1.** Piepho & Möhring broad sense heritability (*H*^2^_PM_) estimated from the joint year model for all traits measured across 2021 and 2022 alongside the signal of selection (*S*) estimated for the same traits using DRIFTSEL. When *S*∼1 this indicates divergent selection between the populations, when S∼0.5 this indicates a neutral pattern (genetic drift), and when S∼0 this indicates stabilising selection between populations.

| Trait | $H^2_{PM}$ | $S$ |
| --- | --- | --- |
| %N | 0.08 | 0.38 |
| %C | 0.13 | 0.39 |
| C:N | 0.05 | 0.41 |
| $\delta^{13}C$ | 0.41 | 0.43 |
| NPQ<sub>ind-amp</sub> | 0.34 | 0.46 |
| NPQ<sub>ind-rate</sub> | 0.39 | 0.36 |
| NPQ<sub>rel-amp</sub> | 0.38 | 0.58 |
| NPQ<sub>rel-rate</sub> | 0.55 | 0.4 |
| NPQ<sub>rel-res</sub> | 0.52 | 0.39 |
| ΦPSII<sub>rec-amp</sub> | 0.38 | 0.6 |
| ΦPSII<sub>rec-rate</sub> | 0.56 | 0.37 |
| ΦPSII<sub>rec-res</sub> | 0.46 | 0.47 |
| NPQ<sub>linear</sub> | 0.59 | 0.34 |
| Maximum NPQ | 0.44 | 0.53 |
| Final NPQ | 0.54 | 0.38 |
| Final ΦPSII | 0.29 | 0.37 |
| DTH | 0.28 | 0.44 |
| Leaf mass | 0.72 | 0.43 |
| Leaf areas | 0.81 | 0.65 |
| SLA | 0.44 | 0.99 |
| V<sub>cmax</sub> | 0.49 | 0.84 |
| J<sub>max</sub> | 0.01 | 1 |
| TPU | 0.31 | 0.36 |
| SL | 0.14 | 0.34 |
| A<sub>sat</sub> | 0.34 | 0.64 |
| g<sub>s</sub> | 0.29 | 0.4 |
| iWUE | 0.01 | 0.59 |

We next estimated the signal of selection (*S*) for all traits using DRIFTSEL (Karhunen & Ovaskainen, 2012). This allowed us to test for the presence of differing evolutionary pressures across the subpopulations for these traits. The average *S* value for all traits was 0.50 which is indicative of a neutral pattern of selection between the subpopulations due to genetic drift (Table 1). However, a few traits did demonstrate strong evidence of divergent selection between the subpopulations, where *S* exceeded 0.95. For example, SLA had a *S* value of 0.99, which may suggest that growth rate is under strong divergent selection across the subpopulations. However, it is also important to note that variation in SLA across the subpopulation may also be a consequence of other determinant factors that influence SLA, such as water availability (Jameel *et al*., 2024) and shade tolerance (Liu *et al*., 2016a). Contrastingly, the *S* value for DTH was moderate (0.44) suggesting that any potential differences in selection for growth rates between the subpopulations may be independent of floral transitioning. Interestingly, the parameters derived from the Rubisco-limited (*V*_cmax_) and electron transport-limited (*J*_max_) portion of the *A*-*C*_i_ curve both demonstrated high S-values. We must apply caution in suggesting that this highlights differential selection for either Rubisco-mediated carboxylation or electron transport since *A*-*c*_i_ curves were only performed on a subset of the total accessions across both years (Supporting Table S1).

We examined correlations across all pairwise trait interactions (Figure 3, Supporting Figures S11-S12). In general, pairwise trait correlations held true across the two common garden experiments (Supporting Figures S10-11), with some exceptions. For example, in 2021, DTH demonstrated a strong positive correlation with *A*_sat_ and *<u>g</u>*_s_ (Supporting Figure S11) suggesting that accessions that were more photosynthetically active tended to be those that had transitioned to flowering later. In 2022, conversely, *A*_sat_ was not associated with DTH (Supporting Figure S12). Additionally, in 2022 we observed multiple significant correlations between δ^13^C and traits that relate to the response of NPQ and ΦPSII to dynamic irradiance that were not present in 2021 (Supporting Figures S10-11). δ^13^C is a useful proxy for WUE integrated over the time in which the carbon forming the tissue was fixed (Leakey *et al*., 2019), consequently these associations may suggest that variation in the kinetics of the light-dependent photosynthetic reactions may have had more of a bearing on variation in WUE in 2022 than in 2021. Consistent with this assertion, we only observed a significant (negative) association between δ^13^C and *<u>g</u>*_s_ in 2021 but not in 2022 (Supporting Figure S11), thereby suggesting that variation in stomatal physiology and behaviour may have been more important in defining variation in WUE in 2021. This suggestion is further supported by the presence of a significant, negative association between *<u>g</u>*_s_ and *iWUE* that was observed in 2021 only (Supporting Figure S11).

**Figure 3.**
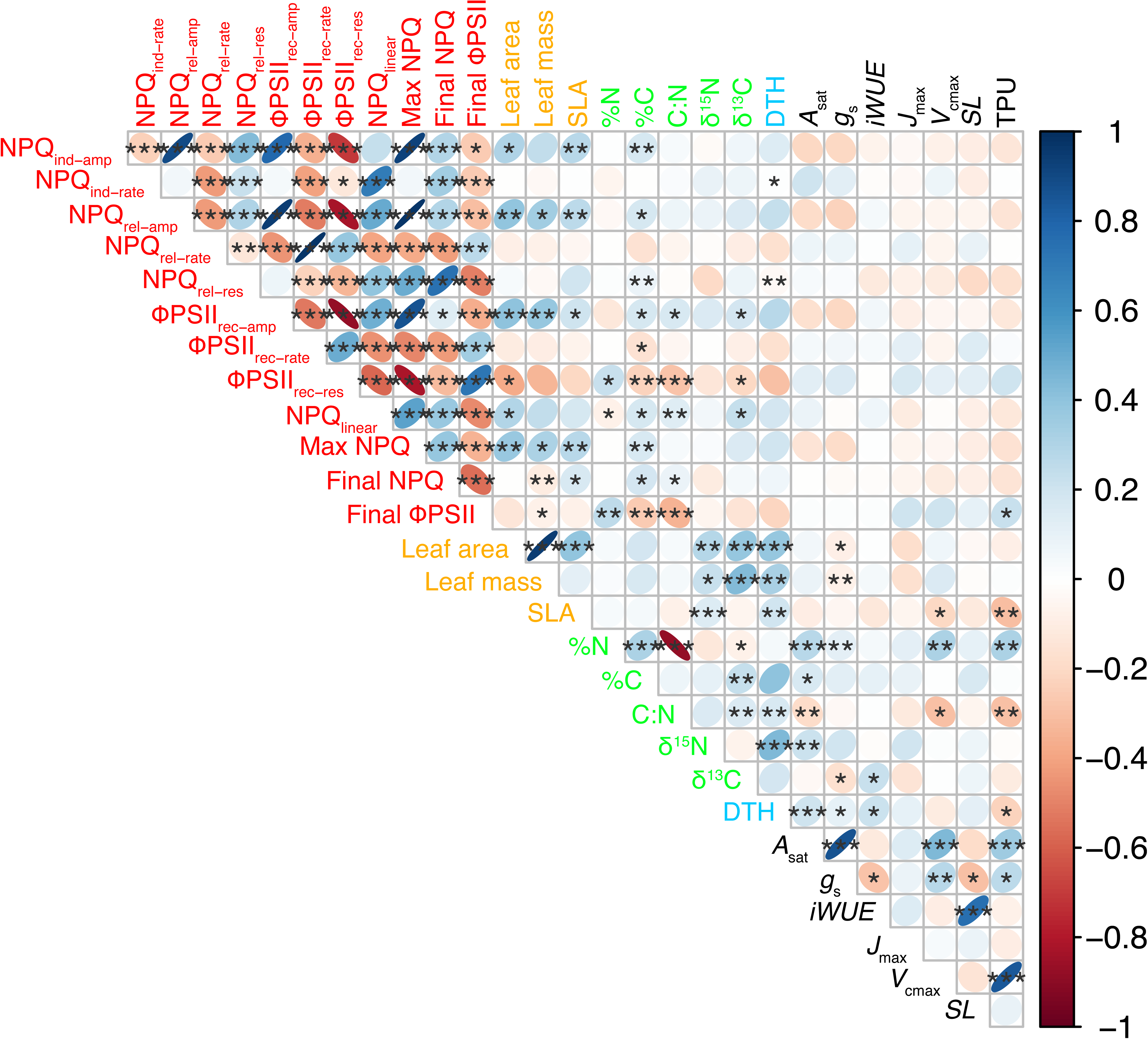
Correlogram showing pairwise trait correlations between traits (BLUEs from the joint model). Significant correlations are highlighted with asterisks at α = 0.05 (*), 0.01 (**), and 0.001 (***). The colour of the ellipses indicated the direction of the correlations. Trait names are coloured according to associated trait groupings. Red: Light dependent reaction-associated traits, Orange: Leaf structure-associated traits, Green: Leaf chemical composition-associated traits, Black: Gas exchange-associated traits

By examining pairwise trait correlations using the joint-year BLUEs we were able to determine phenotypic associations that persisted despite apparent GxE interactions (Figure 1E-H, Supporting Figure S10). Interactions of interest here included the positive correlations between the amplitude of NPQ induction and relaxation with SLA and the amplitude of the recovery of ΦPSII with SLA (Figure 3). These associations may suggest that larger responses of these light dependent reaction-associated parameters to changes in irradiance may be associated with growth rate and/or other determinants of variation in SLA. In terms of photosynthetic capacity, we observed expected associations that highlight the importance of leaf nitrogen content and *g*_s_ as determinants of *A*_sat_ (Figure 3). Further we observed a positive association between *V*_cmax_ and *A*_sat_, but not *J*_max_ and *A*_sat_, which suggests that variation in carboxylation by Rubisco is more important in limiting net CO_2_ assimilation than capacity for RuBP regeneration across this diversity set. We also observed a positive association between *A*_sat_ and TPU as derived from the *A*-*C*_i_ curve (Figure 3), which suggest that more effective export of triose phosphate is linked to higher rates of *A*_sat_ (Lombardozzi *et al*., 2018).

### Observed phenotypic variation is regulated by multiple QTL and shows evidence for local adaptation between subpopulations

For our GWAS, we adopted multiple approaches for detecting SNP-trait associations (See Materials & Methods). Through these various approaches we identified 193 QTL using the joint-year data and 159 QTL using the individual year data. We focused on those that were identified using the joint-year data as these were likely to be more genetically robust. Further, we prioritised *high confidence* QTL as those that were identified using both the joint-year BLUEs and the joint-year BLUPs. In total, we identified 22 high-confidence QTL that were located across all chromosomes except chromosome 6H (Table 2.). Seven of these were associated to DTH, one was identified for SLA, and one was associated to the mass of the penultimate leaf. The remaining 13 QTL were linked to traits associated to light-dependent photosynthetic processes phenotyped via chlorophyll fluorescence. Some of these 13 QTL were non-unique since they were identified for closely related or co-dependent traits, for example the QTL for NPQ_ind-amp_ and maximum NPQ on chromosome 2H.

**Table 2.**
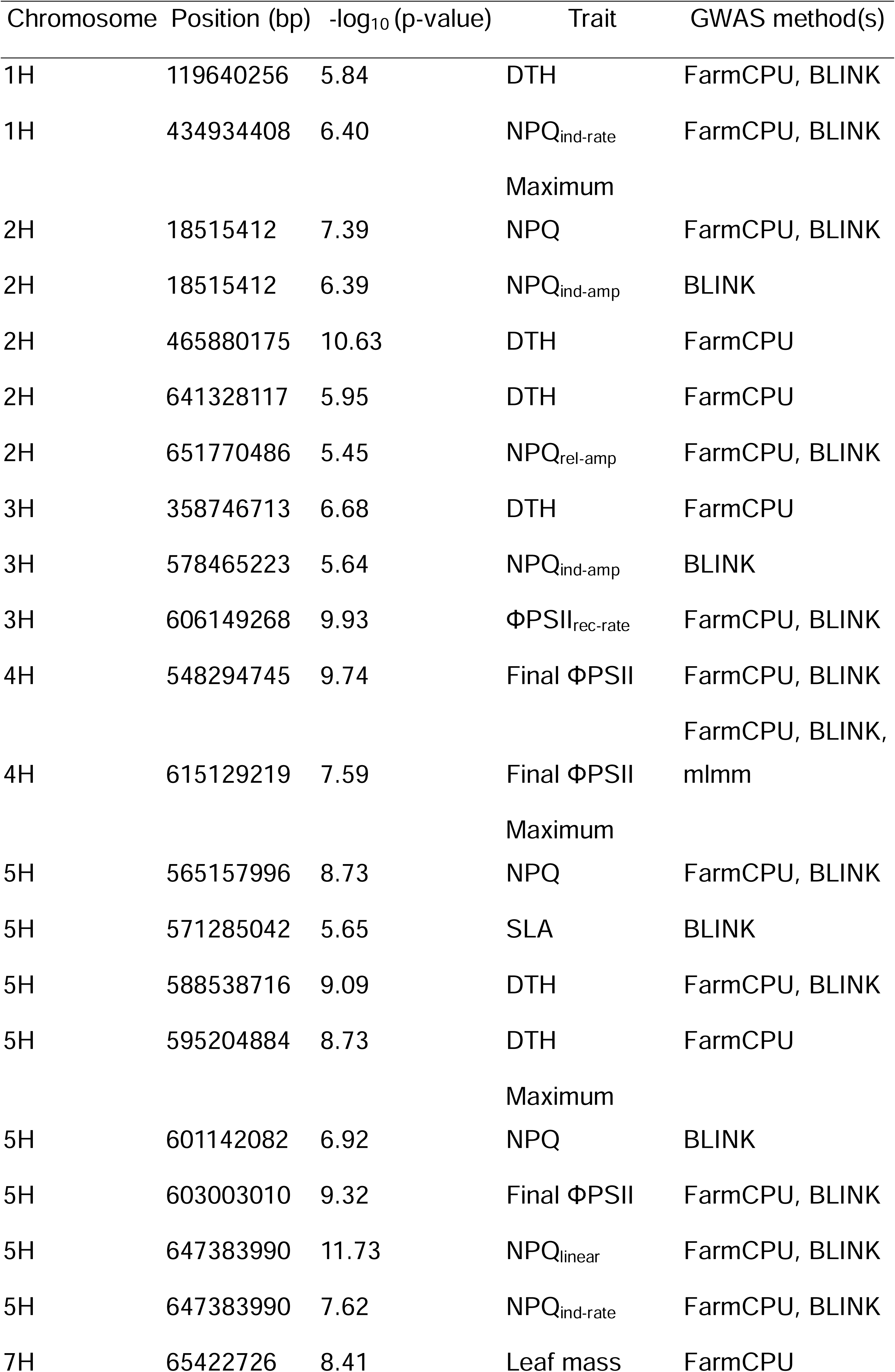
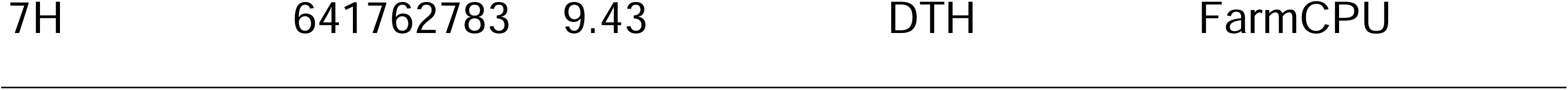
High-confidence quantitative trait loci (QTL) identified. Each row represents a significant SNP-trait association. The chromosome, basepair position, -log10(p-value), associated trait and method of detection are provided.

While the goal of this study was not candidate gene identification, we did explore all the genes within 100kb (upstream and downstream) of these high-confidence QTL. In only one instance did we observe a highly obvious associated candidate gene. This was for the above-described QTL associated to NPQ_ind-amp_ and maximum NPQ on chromosome 2H. Here, we observed just one gene within the 200kb upstream and downstream window, which was ∼49kb away from the SNP defining this locus and annotated as Rubisco small subunit (*rbcS*; HORVU.MOREX.r3.2HG0104730). *rbcS* mutants are known to have perturbed NPQ light responses and reduced maximum NPQ (Atkinson *et al*., 2017), thereby highlighting the plausibility of *rbcS* as a candidate gene here although further validation would be needed.

For all the high confidence QTLs we tested for differences in the allelic frequency of the SNPs associated to those QTL between the identified subpopulations following the approach of Fustier *et al*., (2019). Here, we identified three QTLs/SNPs both allelic frequencies and trait variation for which the QTLs had been identified were significantly different between the subpopulations (Figure 4).

**Figure 4.**
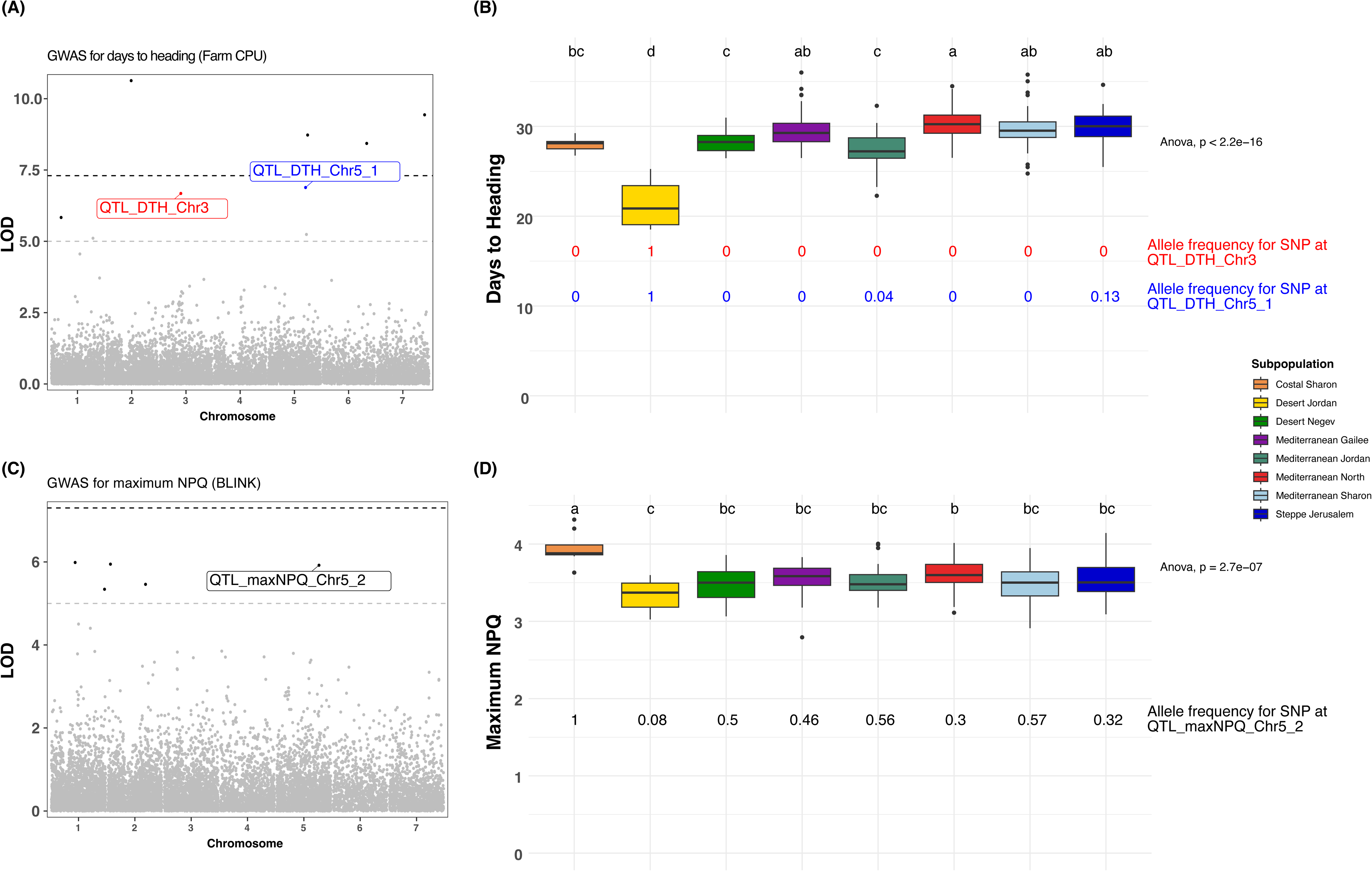
GWAS and follow up analyses of allelic frequency of significant SNPs. (A) GWAS for days to heading (DTH). SNPs where allelic frequency across the subpopulations is different and mirrors associated trait variation are highlighted, with those differences being demonstrated in panel B. (C) GWAS for maximum non-photochemical quenching (NPQ). A SNP whose allelic frequency across the subpopulations is different and mirrors associated trait variation is highlighted, with those differences being demonstrated in panel D.

The QTLs identified for DTH on chromosome 3H and 5H (1^st^ QTL) demonstrated distinct allelic frequencies with respect to the Desert Jordan subpopulation, which also demonstrated markedly earlier flowering than the remaining subpopulations (Figure 4A-B). Given the significantly reduced precipitation that characterises the region of origin for this subpopulation (Figure 1D), it could be suggested that there has been active selection on the alternative allele at these two QTL in the Desert Jordan subpopulation to promote early flowering as a drought adaptive mechanism.

The second QTL identified for maximum NPQ on chromosome 5H had significantly higher allelic frequency in the Coastal Sharon subpopulation which also demonstrated significantly higher maximum NPQ compared to the remaining subpopulations (Figure 4C-D). Coastal areas tend to receive more variable cloud cover and enhanced cloudy environments have been shown to elicit higher maximum NPQ in Arabidopsis ecotypes due to lack of adaptation to high light environment (Rungrat *et al*., 2019). Thus, it is plausible to suggest the alternative allele at this QTL may be linked to an NPQ-derived adaptation to the high-light, therefore it is selected for in the non-Coastal Sharon accessions.

In order to further understand physiological mechanisms that may underpin differential adaptation across these subpopulations, we compared both absolute trait variation as well as variation in trait plasticity across the subpopulations (between the 2021 and 2022 growing season; see Materials and Methods for details). Through this approach, we observed that the desert subpopulations were most commonly the outliers relative to the other subpopulations for both plasticity and absolute metrics (Figure 5; Supporting Figure S13-S14). For example, the Desert Jordan subpopulation demonstrated significantly reduced SLA compared to all other subpopulations except the Mediterranean Sharon subpopulation (Figure 5A). Curiously, the only subpopulation to demonstrate significantly different δ^13^C compared to other subpopulations was the Desert Negev subpopulation (Figure 5B), which tended to show the most negative values which would suggest reduced water-use efficiency. It is important to note that this may also be a result of differential post-photosynthetic fractionation in the Desert Negev subpopulation. The Desert Jordan subpopulation displayed by far the fastest flowering time, since DTH was significantly reduced in this subpopulation compared to all other subpopulations (Figure 5C). We did not detect any significant differences between the subpopulations for *g*_s_, however it is notable that the Coastal Sharon subpopulation, which is characterised by the highest average monthly precipitation (Figure 1D), had by far the highest average *g*_s_ (Figure 5D).

**Figure 5.**
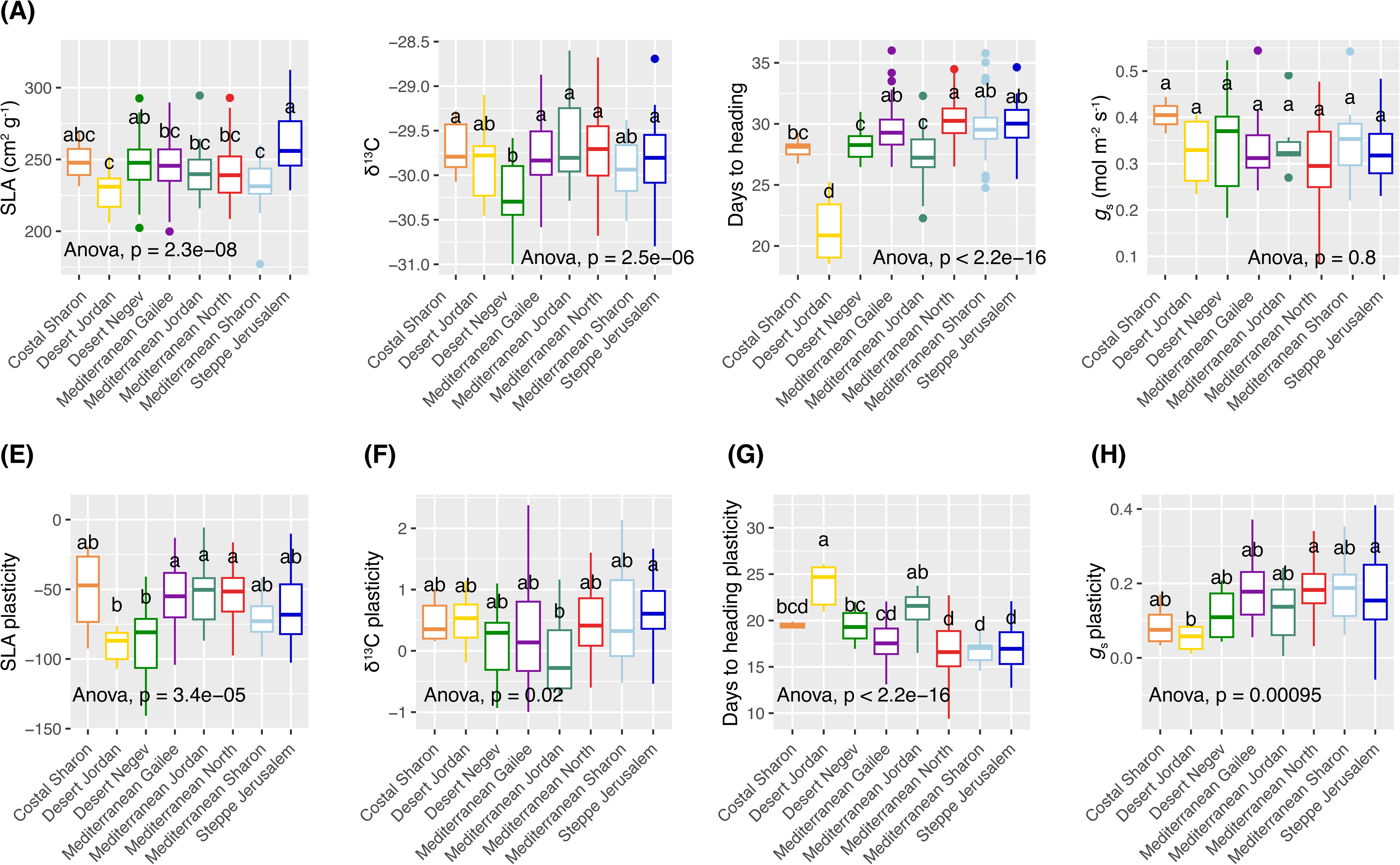
Trait variation across the subpopulations for (A) SLA, (B) carbon isotope composition (δ^13^C), (C) stomatal conductance (*g*_s_), and (D) days to heading (DTH). Variation in plasticity between 2021 and 2022 across the subpopulations for (E) SLA, (F) δ^13^C, (G) *g*_s_, and (H) DTH. P-values associated to one-way ANOVA comparison of means tests are inset and associated post-hoc classifications of the subpopulations are indicated by the letters above the boxplots.

With respect to differences in plasticity, the two Desert subpopulations showed the greatest change in SLA between the two growing season, where the variation in plasticity across these two subpopulations was significantly more negative (indicating a shift towards higher SLA in 2022) than three of the four Mediterranean subpopulations (Figure 5E). Variation in plasticity for δ^13^C was relatively consistent across the subpopulations, with only the Mediterranean Jordan and Steppe Jerusalem subpopulations showing a significant difference (Figure 5F). Conversely, plasticity in DTH was much more variable with four post-hoc groups identified. Here, the Desert Jordan subpopulation demonstrated the greatest plasticity (indicating a shift towards earlier flowering in 2022) with the Steppe Jerusalem subpopulation showing the least plasticity between the growing seasons (Figure 5G). These two subpopulations were also the only two to demonstrate significantly different variation in *g*_s_ plasticity, with the Desert Jordan subpopulation showing relatively low plasticity and the Steppe Jerusalem showing comparatively higher *g*_s_ plasticity (Figure 5H).

To further test the capacity for GxE and its role in adaptation to environments across distinct subpopulations, we selected three accessions each from the Desert Jordan and Steppe Jerusalem subpopulations based on PCA (Supporting Figure S3). These six accessions were grown under two distinct and consistently maintained water availabilities (See Materials and Methods). At the end of this experiment, we measured total leaf area as a proxy of above ground biomass (Figure 6). Plants grown under reduced water availability (40% rSWC) demonstrated reduced total leaf area (Figure 6, Supporting Table S5). This was consistent across the two subpopulations, although some genotypes demonstrated stronger reductions than others. Whilst the observation was not particularly surprising, we do note that the percentage decline was much more extreme across the Steppe Jerusalem genotypes (77.74%) relative to the Desert Jordan genotypes (54.32%).

**Figure 6.**
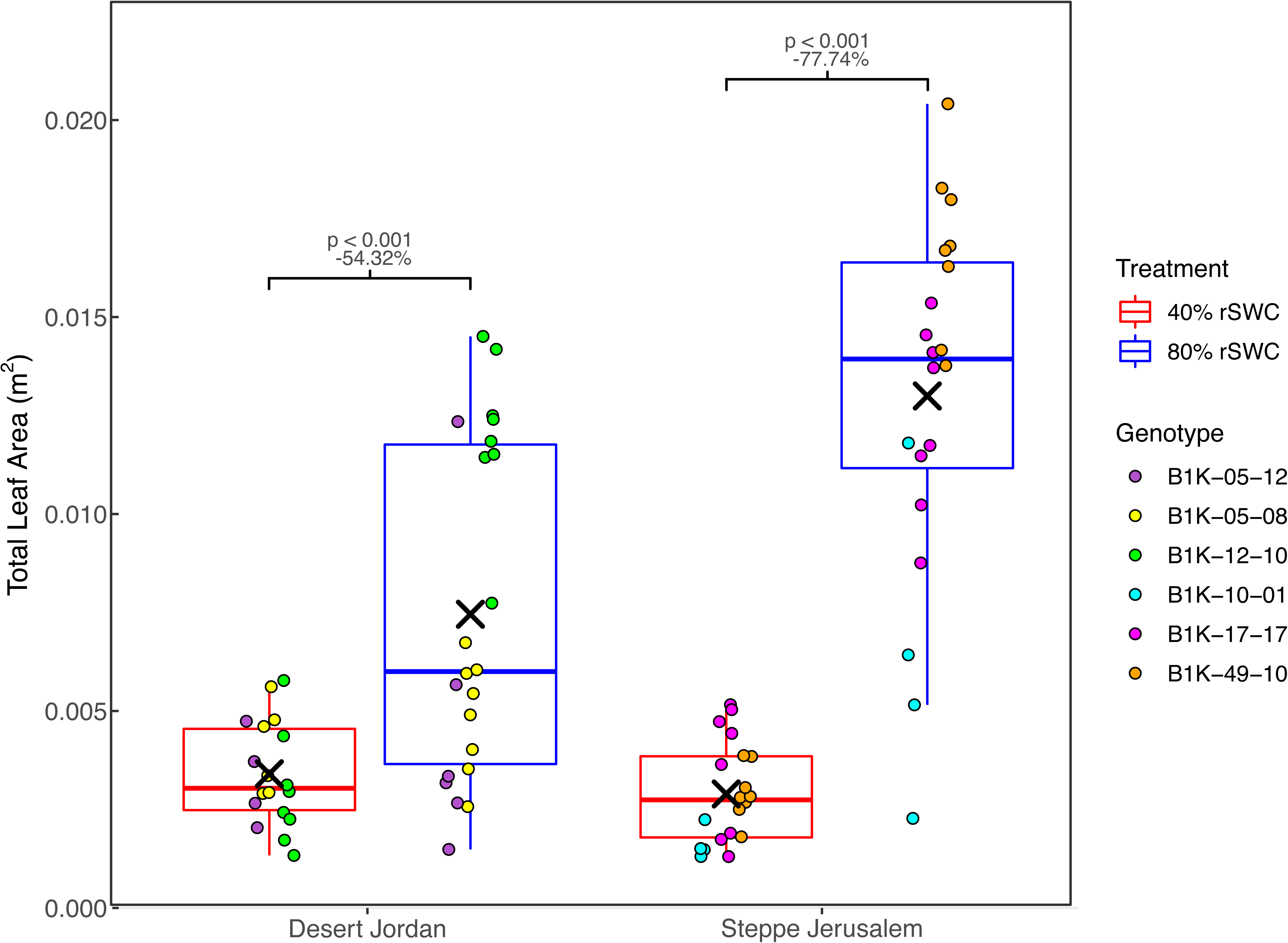
Total leaf area of the six accessions included in the detailed experiment that were subjected to two distinct watering regimes to maintain relative soil water content (rSWC) at 40% and 80%. The percentage change of the mean for each accession between 80% and 40% rSWC is inset within each sub-panel alongside the associated p-value obtained from post-hoc Tukey tests applied to the associated two-way ANOVA

During the above-described water availability experiment we profiled leaf-level gas exchange and performed *A*-*C*_i_ response curves in order to generate data that may explain differences in biomass accumulation. Here we observed that *V*_cmax_ and *J*_max_ were significantly reduced across the Steppe Jerusalem accessions when grown at 40% rSWC compared to 80% rSWC (Figure 7A-B; Supporting Table S5). However, this effect was not mirrored by the Desert Jordan accessions, where neither *V*_cmax_ nor *J*_max_ were significantly reduced when associated accessions were grown at 40% rSWC compared to 80% rSWC (Figure 7A-B; Supporting Table S5).

**Figure 7.**
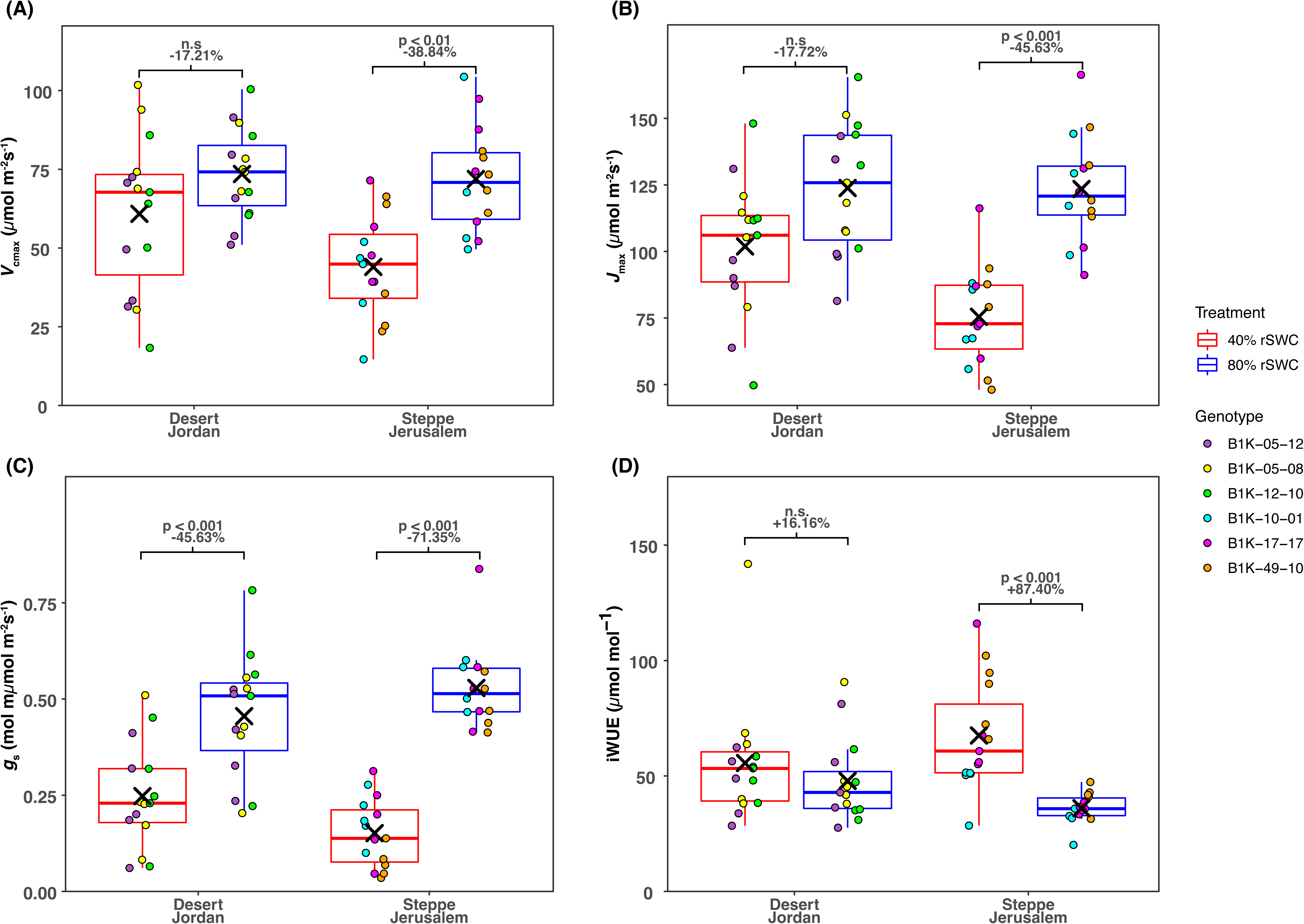
Photosynthesis-associated traits of the six accessions included in the detailed experiment that were subjected to two distinct watering regimes to maintain relative soil water content (rSWC) at 40% and 80%. (A) Maximum rate of carboxylation by rubisco (*V*_cmax_). (B) Maximum rate of electron transport for RuBP regeneration (*J*_max_). (C) Stomatal conductance (*g*_s_). (D) Intrinsic water-use efficiency (*iWUE*). The percentage change of the mean for each accession between 80% and 40% rSWC is inset within each sub-panel alongside the associated p-value obtained from post-hoc Tukey tests applied to the associated two-way ANOVA

The differences in GxE for photosynthetic capacity across the two subpopulations may in part be explained by effects on *g*_s_ (Figure 7C; Supporting Table S5). We observed that *g*_s_ was significantly reduced for both subpopulations when grown at 40% relative to 80% rSWC, however this reduction was much greater (71.35%) for the Steppe Jerusalem subpopulation than the Desert Jordan subpopulation (45.63%). This suggests that the Desert Jordan accessions are able to maintain normal photosynthetic capacity under reduced water availability despite reduced *g*_s_.

Notwithstanding the significant decline in *g*_s_, *iWUE* of the Desert Jordan accessions was not significantly different between the two water availability treatments (Figure 7D, Supporting Table S5), which is further reflective of the adaptive capacity of the Desert Jordan accessions and suggests that despite restricting water loss, there is no modulation between the amount of carbon gained per unit of water lost under the tested drought scenario. Conversely, *iWUE* of the Steppe Jerusalem accessions was significantly increased when plants were grown under reduced water availability (Figure 7D; Supporting Table S5), suggesting that accessions from this subpopulation may need to fine-tune this trade off relative to how they perform under optimal resource availability.

## Discussion

Exploiting natural variation in photosynthesis and introgressing new genetic diversity from CWRs into elite cultivars are two strategies toward crop improvement that have gained significant attention in recent years (Zhang *et al*., 2017; Prohens *et al*., 2017; Faralli & Lawson, 2020; Theeuwen *et al*., 2022; Renzi *et al*., 2022). Combining these approaches could have enormous potential for increasing photosynthesis in crops. However, we presently have a very limited understanding as to the extent with which photosynthesis varies intraspecifically in any CWR of interest. With this study, we address this knolwdge gap for the first time through a comprehensive assessment of natural variation of multiple photosynthesis-associated traits across wild barley. Through phenotyping 320 accessions across two growing seasons (270 common across both years) we showcase a wide toolbox of photosynthetic diversity for future barley improvement efforts. Our results suggest that SLA and biochemical capacity for CO2 assimilation are subject to divergent selection. Further, three QTLs identified for DTH and NPQ show allelic frequency distributions between subpopulations that are consistent with a role in local adaptation. In the following paragraphs, we provide context on how this natural variation could be harnessed for crop improvement in the context of climatic change.

Since the genetic structure of many of the accessions incorporated in our study had not previously been quantified, we performed associated analyses as an important first step for this study. We identified eight distinct subpopulations (Figure 1B-C) that largely aligned with the recent work of Chang *et al*., (2022), consistent with the presence of 164 common accessions between the two (Supplemental Table S1). We obtained climatic data from the point of collection of all accessions and identified distinct differences in historical water availability and growing temperatures (Figure 1D; Supporting Figure S6). These environmental parameters are well-known to have strong and multifaceted effects on photosynthesis (Chaves *et al*., 2009; Yamori *et al*., 2014) and are critical in determining local adaptation (Stebbins, 1952). Consequently, the B1K accessions incorporated in this study represent an interesting panel through which to quantify variation across the *photosynthome* of a CWR, but also for understanding how this variation could be shaped by local adaptation to distinct abiotic factors.

In our study, we focused our measurements on the penultimate leaf, which is important for defining grain filling rate and grain quality in barley (Shirdelmoghanloo *et al*., 2022). We measured the SLA of this leaf, which is defined by the ratio of leaf area and dried leaf mass. It is important trait for defining investment of resources and plant growth (Lambers & Poorter, 1992). Moreover, it has been utilised in barley breeding contexts, where it has been demonstrated to be a reliable physical marker of grain yield (Alqudah & Schnurbusch, 2015) and vigour (Rebetzke *et al*., 2004). Consequently, we utilise SLA as a proxy for biomass accumulation in our study. In alignment with previous observations in domesticated barley (Rebetzke *et al*., 2004; Alqudah & Schnurbusch, 2015), we observed a strong genetic component to SLA in wild barley (Table 1). Moreover, alongside DTH (Supporting Figure S10), SLA and its constituents showed the greatest similarity in variation across the growing seasons (Figure 2, Supporting Figure S10), suggesting reduced GxE compared to the photo-physiological traits measured. In addition, SLA demonstrated a very high *S* value (Table 1), suggesting that there is differential selection for SLA across the defined subpopulations (Figure 1). In general, SLA increases with resource availability in the short term, however long-term adaptation to unfavourable habitats can also result in SLA increasing over evolutionary time despite lack of resources (Liu *et al*., 2023). In this context, we note that accessions from the two desert environments demonstrate a significant difference in subpopulation-wide SLA variation (Figure 5A). Here, the Desert Jordan subpopulation demonstrated the lowest SLA, which is line with an expected adaptation to limited water availability (Scheepens *et al*., 2010; Wilcox *et al*., 2021). However, variation in SLA across the Desert Negev subpopulation was similar to the majority of other subpopulations, suggesting that these accessions may have undergone longer term adaptation to reduced precipitation and/or there is a strong influence on SLA coming from other environmental parameters, e.g., light availability (Liu *et al*., 2016a).

In barley, like many other species, there is a strong association between light availability and SLA (Gunn *et al*., 1999). To this end, we observed significant positive associations between SLA and traits that define the response of NPQ and ΦPSII to light (Figure 3). Specifically, our results highlight that greater changes in the induction and relaxation of NPQ and recovery of ΦPSII are linked to greater SLA, and potentially therefore biomass accumulation. A similar observation was also made recently by Cowling *et al*., (2022) who showed that NPQ dynamics were linked to biomass accumulation across >150 diverse African rice landraces. Interestingly, Cowling *et al*., (2022) also demonstrated that photophysiological diversity as a whole was linked to ecological niches, but they did not specifically test any particular photosynthesis-related traits. In our study, we also observed differences between subpopulations for NPQ and ΦPSII dynamics (Supporting Figures S13), further highlighting how dynamic photoprotection and photosynthesis may be linked to local adaptation. Moreover, through our GWAS we detected many marker-trait associations for these NPQ- and ΦPSII-related traits (Table 2). In one case, a very obvious candidate gene was in close proximity to an identified marker. Here, *rbcS* was linked to genetic variation in the amplitude of change in NPQ upon transitioning from high light to darkness, which follows previous observations in Arabidopsis mutants (Atkinson *et al*., 2017). This could relate to synchronisation between the demand for ATP for CO2 fixation and acidification of the thylakoid lumen, which triggers NPQ (Ramakers *et al*., 2025). In another instance, we observed a link between the frequency of alleles of a marker significantly associated to maximum NPQ and subpopulation variation in maximum NPQ (Figure 4C-D). Here, the alternative allele appeared fixed in the Coastal Sharon subpopulation, which also demonstrated significantly higher maximum NPQ compared to any other subpopulation. In general, coastal environments tend to be characterised by more cloud cover, which likely places significant pressure to adapt the response of photoprotection to dynamic light. A study focusing on diversity in Arabidopsis showed that growing plants under coastal (low light) and inland (high light) environments results in differing NPQ responses to light and darkness, where plants growing under coastal conditions have higher maximum NPQ (Rungrat *et al*., 2019). This may initially appear counterintuitive; however, it may be a result of inland grown plants being adapted to initiate appropriate physiological responses to deal with stressful high-light conditions since they are more frequently exposed to them than coastal plants. Thus, in our study, it may be the case that Coastal Sharon accessions are not exposed to strong selection pressure to adapt to a high-light environment, thus there is no variation in the identified allele and maximum NPQ is not as well constrained as it is in the other subpopulations, which may have consequences for photosynthesis and productivity.

Similar observations were made for two markers significantly associated to variation in DTH (Figure 4A-B). Here, the alternative allele at these two markers was dominant in the Desert Jordan subpopulation, which also had the shortest DTH of any subpopulation. This is plausibly a result of evolution of a drought escape strategy in the Desert Jordan subpopulation, which may be associated to genetic variation at or close to these markers. This strategy describes how natural populations evolve to complete their lifecycle more rapidly to ensure that they don’t succumb to a drought event (Reviewed by: Kooyers, 2015). Indeed, previous common garden experiments have shown that populations from xeric environments tend to flower earlier (e.g., Knight *et al*., 2006; Lowry *et al*., 2015). As with the SLA observation described above, this trend does not hold true with the Desert Negev subpopulation, suggesting that this environment is not as stressful in term of water availability and/or these accessions have evolved alternative drought avoidance mechanisms. In this instance, drought avoidance is unlikely to be related to differences in leaf-level gas exchange, since neither *g*_s_ nor δ^13^C were observed to be tailored to a water limited environment in the Desert Negev subpopulation (Figure 5B, D). It is important to note the drought escape strategy does not necessarily evolve as a constitutive response that occurs independently of environmental cues. It can also evolve as a heritable plastic response (Riboni *et al*., 2013). To this end, we also note that the Desert Jordan subpopulation displayed the greatest plasticity for DTH across the two growing seasons, suggesting that accessions from this subpopulation are well adapted to persistently and/or sporadically dry environments. Further, these accessions do not appear to suffer a reduction in photosynthetic assimilation when grown under water replete conditions (Supplemental Figure S13) such as those that characterise the common garden experiments performed in this study. This last point is important as it is key that drought tolerant crops are able to additionally perform well in situations that do not occasion drought tolerance (Blum, 2005).

With respect to *A*_sat_, we observed a substantial amount of variation across the diversity panel (Figure 2B) that was reasonably consistent across the two growing seasons (Figure 2F) despite a strong contribution from the environment in determining the variation (Table 1). In general, the heritability estimates of photosynthesis traits estimated from gas exchange in this study relative to the study of domesticated barley outlined by Gao *et al*., (2024) are reduced. However, it is worth noting that the latter performed these measurements using plant material grown in controlled environments, thereby limiting environmental noise. In addition, domesticated barley has been specifically bred for stability (Kraakman *et al*., 2004), which is not the case for wild species, where lack of phenotypic robustness is often advantageous (Lachowiec *et al*., 2016). These considerations may help explain this particular discrepancy between the two studies. Gao *et al*., (2024) observed a mean *A*_sat_ across their studied barley varieties of 17.2 μmol CO_2_ m^-2^ s^-1^, and a maximum of 19.7 μmol CO_2_ m^-2^ s^-1^. In our study, the average value for *A*_sat_ from the joint-year BLUEs was 20.19 μmol CO_2_ m^-2^ s^-1^ and some accessions demonstrated rates exceeding 30 μmol CO_2_ m^-2^ s^-1^ (Figure 1B). The *A*_sat_ data from our 2022 growing season was more comparable to Gao *et al*., (2024), however there were still some accessions with much higher rates of *A_s_*_at_. Moreover, average *A_s_*_at_ in 2021 was 23.31 μmol CO_2_ m^-2^ s^-1^ (Figure 1B). In general, these results showcase that there is a wealth of variation in *A*_sat_ in wild barley that is indicative of photosynthetic rates greater than those measured in demonstrated in domesticated barley. Our work has made a start to better understand the genetics underpinning this, but this may require further experimentation of derived crosses in controlled environments to maximise heritability. This could serve as the foundation for a *pre-breeding* approach to introgress wild photosynthesis-promoting alleles into barley, since the two species are inter-fertile (Ellis *et al*., 2000). Notably, a recent multi-parent double haploid population that includes donation from ten of the B1K accessions (Bodenheimer *et al*. *In prep*) could bridge the utilisation of the QTL identified here to pre-breeding and field trials.

Whilst we were unable to identify QTL associated with *A*_sat_ variation, we were able to assess patterns of variation that help to better understand the basis for this variation. For example, a very strong, positive association was observed with *g*_s_ (Figure 3). This follows the well-characterised trend that enhancing photosynthesis in a C_3_ species typically requires concurrent increases in *g_s_* (Leakey *et al*., 2019). This clearly has implications for WUE; however, we note that the negative association between *g*_s_ and *iWUE* is not as strong as the *g*_s_-*A*_sat_ association. This implies that there are some accessions that can operate at high rates of *A*_sat_ with low or moderate *g*_s_, which would be ideal for breeding barley to drought-prone environments (Rebetzke *et al*., 2004). Indeed, our experiment with six selected contrasting accessions (discussed further below) highlights the trait combinations that could be selected for. Another commonly observed association we detected was the positive association between leaf nitrogen content and *A*_sat_ (Figure 3), which reflects the investment in nitrogen of the Calvin Cycle and thylakoid proteins (Evans & Clarke, 2019). We also observed a significant positive association between δ^15^N and *A*_sat_ (Figure 3). In plants, the physiological mechanisms that influence δ^15^N are not as well understood as those that influence δ^13^C. However, there is some evidence that carbon metabolism and nitrogen assimilation, alongside the isotopic composition of the external nitrogen source, are important in determining δ^15^N (Evans, 2001; Kalcsits *et al*., 2014). Intriguingly, some of the early work to understand genotypic differences in δ^15^N actually comes from wild barley (Handley *et al*., 1997; Robinson *et al*., 2000), where genotypic variation in δ^15^N was linked to abiotic stress tolerance (drought and N-starvation) and also the capacity to retain N, which would have implications for photosynthesis. This may also reflect the association between leaf δ^15^N and DTH (Figure 3), since floral transitioning is typically associated with the remobilisation of nitrogen and the onset of senescence (Distelfeld *et al*., 2014). The importance of leaf nitrogen is further reflected in the positive correlations between *V*_cmax_ and %N and *A*_sat_, which reflects the investment of N in Rubisco (Luo *et al*., 2021) and confirms that Rubisco activity is key for photosynthetic performance in barley.

As a final component to this study, we sought to understand how differences in photophysiological plasticity may contribute to adaptation in wild barley. To this end we focused on accessions from the Desert Jordan and the Steppe Jerusalem subpopulations as the only two groups to show differences in *g*_s_ plasticity (Figure 5D). Plants grown under reduced water availability showed reduced total leaf area across both subpopulations (Figure 6). This is a fairly-well characterised response to water deficits in plants and has also been observed recently in domesticated barley (Moualeu-Ngangué *et al*., 2020). In general, this response reflects (a) an attempt by the plant to reduce surface area for transpiration; and/or (b) a reallocation of resources to promote root growth for water acquisition (Chaves *et al*., 2009). Whilst both subpopulations showed declining leaf area when grown under reduced water availability, the extent of this plasticity was much reduced in the Desert Jordan subpopulation. Moreover, this reduction was primarily driven by a single accession (B1K-12-10). This observation is in line with a recent study that focused on a wild C_4_ grass, *Bouteloua gracilis*, where the authors observed that populations from more arid environments tended to be smaller whilst demonstrating reduced plasticity to water limitation (Bushey *et al*., 2023). The relative maintenance in vegetative biomass accumulation in Desert Jordan accessions might be related to their ability to maintain photosynthetic capacity under reduced water availability (Figure 7A-B). Indeed, recent work on wild relatives of wheat has shown that maintaining photosynthesis under drought stress is key to maintaining biomass accumulation and that this capacity is limited in commercial wheat varieties (Mahmood *et al*., 2023). Given the link between *g*_s_ and photosynthetic capacity (Paillassa *et al*., 2020), we were surprised to see that *g*_s_ significantly declined in the Desert Jordan accessions as well as the Steppe Jerusalem accessions (Figure 7C). Again, however, the magnitude of this response was far greater for the Steppe Jerusalem accessions. In light of this observation and given the previously described importance of *g*_s_ for positively defining photosynthesis in wild barley (Figure 3), it suggests that these Steppe Jerusalem accessions may have evolved mechanisms that allow conserved water use by reducing stomatal opening with limited negative effects on photosynthesis. This means they do not have to elicit a significant *iWUE* response, in contrast to the Steppe Jerusalem accessions (Figure 7D). In general, this highlights a degree of uncoupling between *g*_s_ and photosynthesis in the Desert Jordan accessions, which is unusual for a C_3_ species but highly attractive from the standpoint of improving the productivity of crops in water limited environments (Condon *et al*., 2004; Blum, 2005; Leakey *et al*., 2019). Therefore, these accessions warrant further investigation given the importance of improving barley drought tolerance in the future (Elakhdar *et al*., 2022).

## Acknowledgements

This work was supported by the European Union’s Horizon2020 research and innovation 657 programme (No.862201) project CAPITALISE. We thank Haidee Philpott and Dr Tally Wright for consultation on experimental design and statistical analyses. We are grateful to Dr Jenia Binenbaum for assistance with naming the subpopulations.

## Supporting Information

Supporting Figure S1. Common garden sites

Supporting Figure S2. Phenotyping flow chart

Supporting Figure S3. PCA biplot

Supporting Figure S4. Temp and RH in glasshouse

Supporting Figure S5. PAR in the glasshouse

Supporting Figure S6. Boxplots showing mean monthly temperature for site of origin of all subpopulations

Supporting Figure S7. (A) Maximum and minimum temperature during 2021 common garden experiment. (B) Maximum and minimum temperature during 2022 common garden experiment. (C-D) Differences in temperature between 2021 and 2022.

Supporting Figure S8. (A-B) Daily water input (precipitation and irrigation) during 2021 and 2022 common garden experiment. (C) Average daily water input in 2021 and 2022.

Supporting Figure S9. Density plots showing trait variation for traits not shown in Figure 2

Supporting Figure S10. Scatter plots showing correlations between years for traits not shown in Figure 2.

Supporting Figure S11. Pairwise trait correlations for 2021

Supporting Figure S12. Pairwise trait correlations for 2022

Supporting Figure S13. Boxplots for each subpopulation for traits not shown in Figure 5

Extra supporting Figure to fit in somewhere: boxplot of average monthly temperature across the subpopulations

Supporting Figure S14. GxE plots for traits not shown in Figure 5

Supporting Table S1. List of accessions comprising both common garden experiments

Supporting Table S2. List of agronomic inputs for both common garden experiments

Supporting Table S3. Variance associated with each term for all models

Supporting Table S4. Heritabilities derived from year-specific models

Supporting Table S5. Results from two-way ANOVAs performed using glasshouse experiment data

