## Supplementary figures and images for "Extensive photophysiological variation in wild barley is linked to environmental origin"

### Supplemental Figure S1

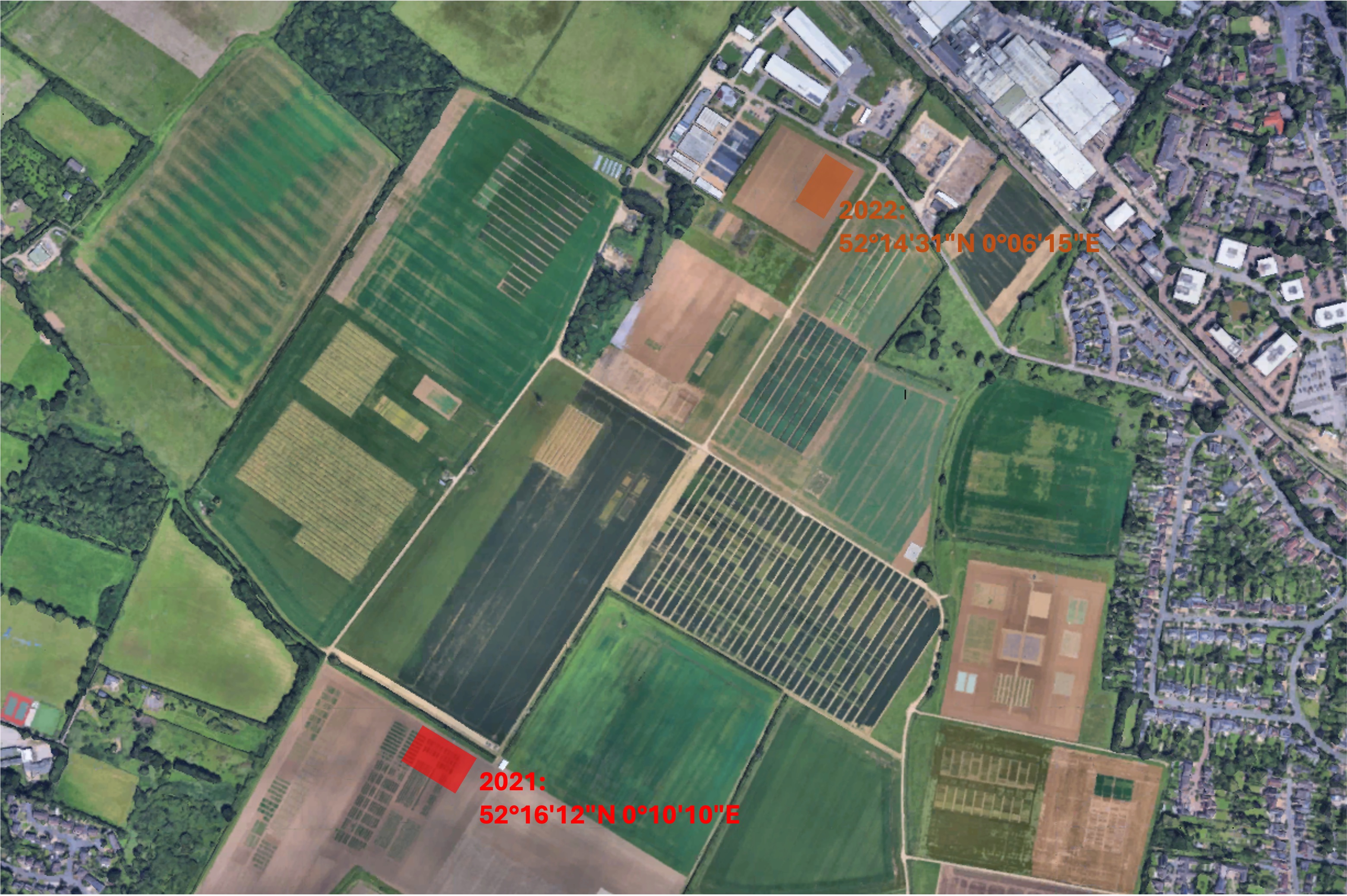

### Supplemental Figure S2

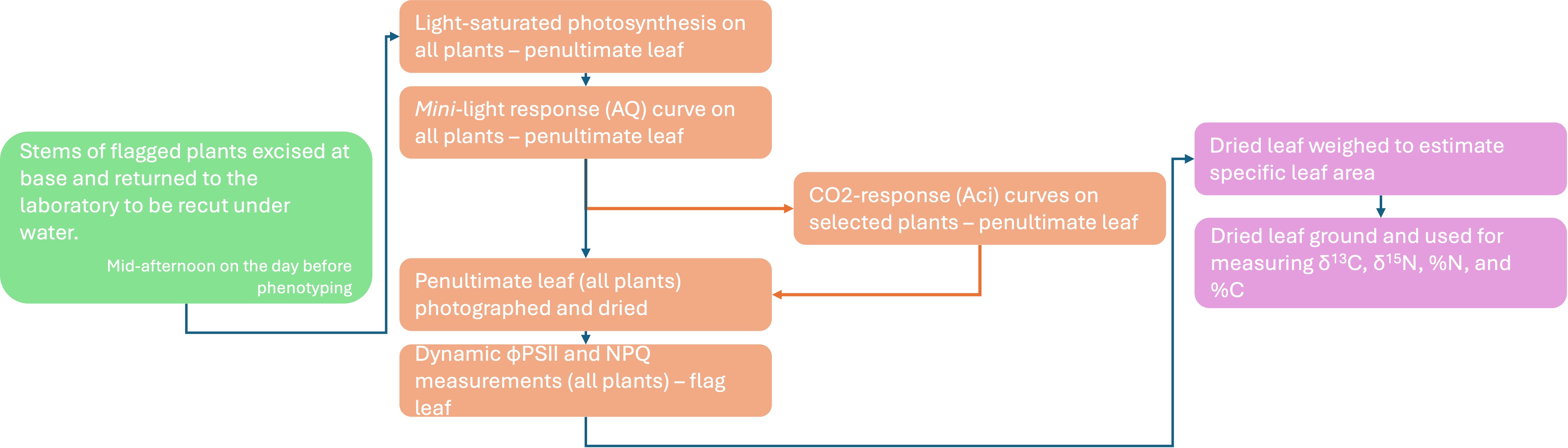

### Supplemental Figure S3

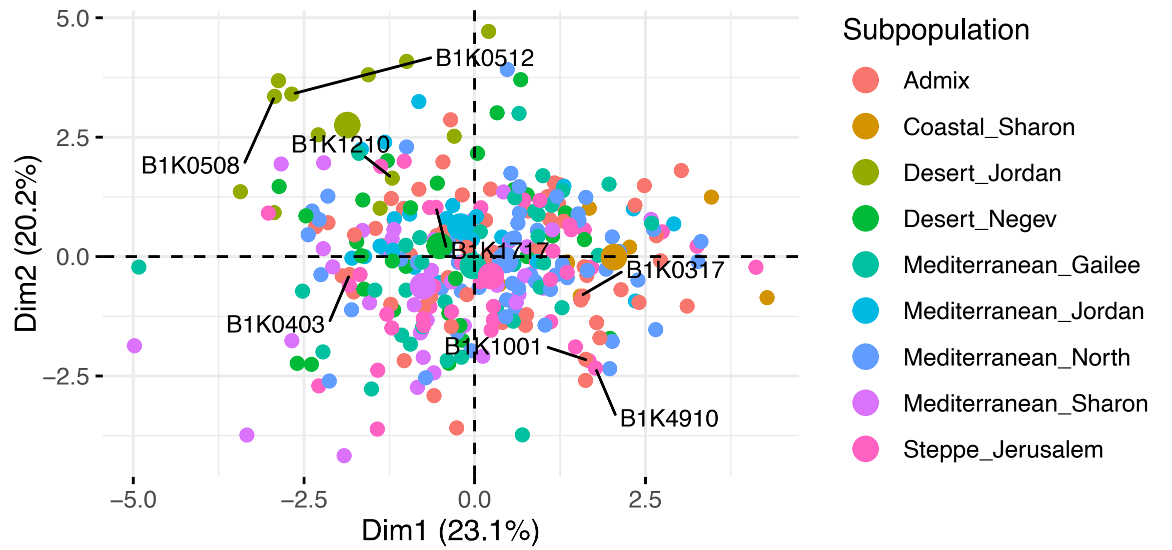

### Supplemental Figure S4

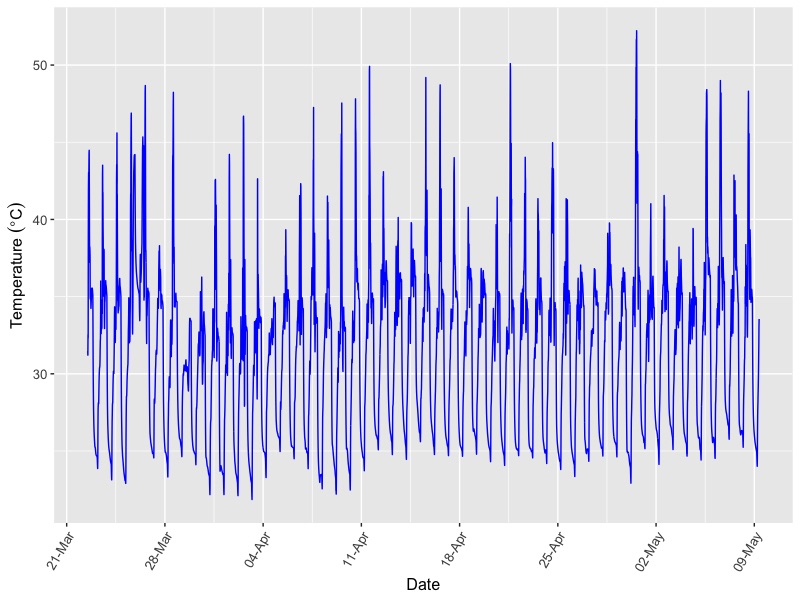

### Supplemental Figure S5

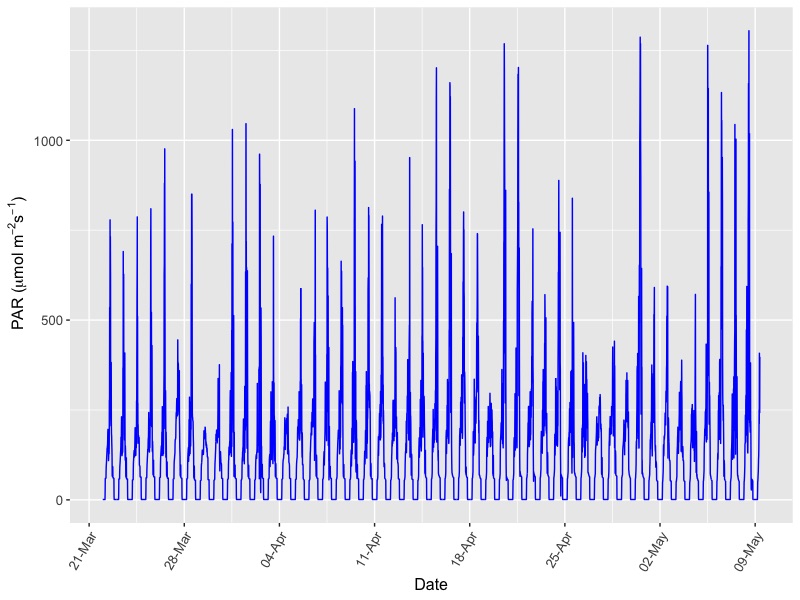

### Supplemental Figure S6

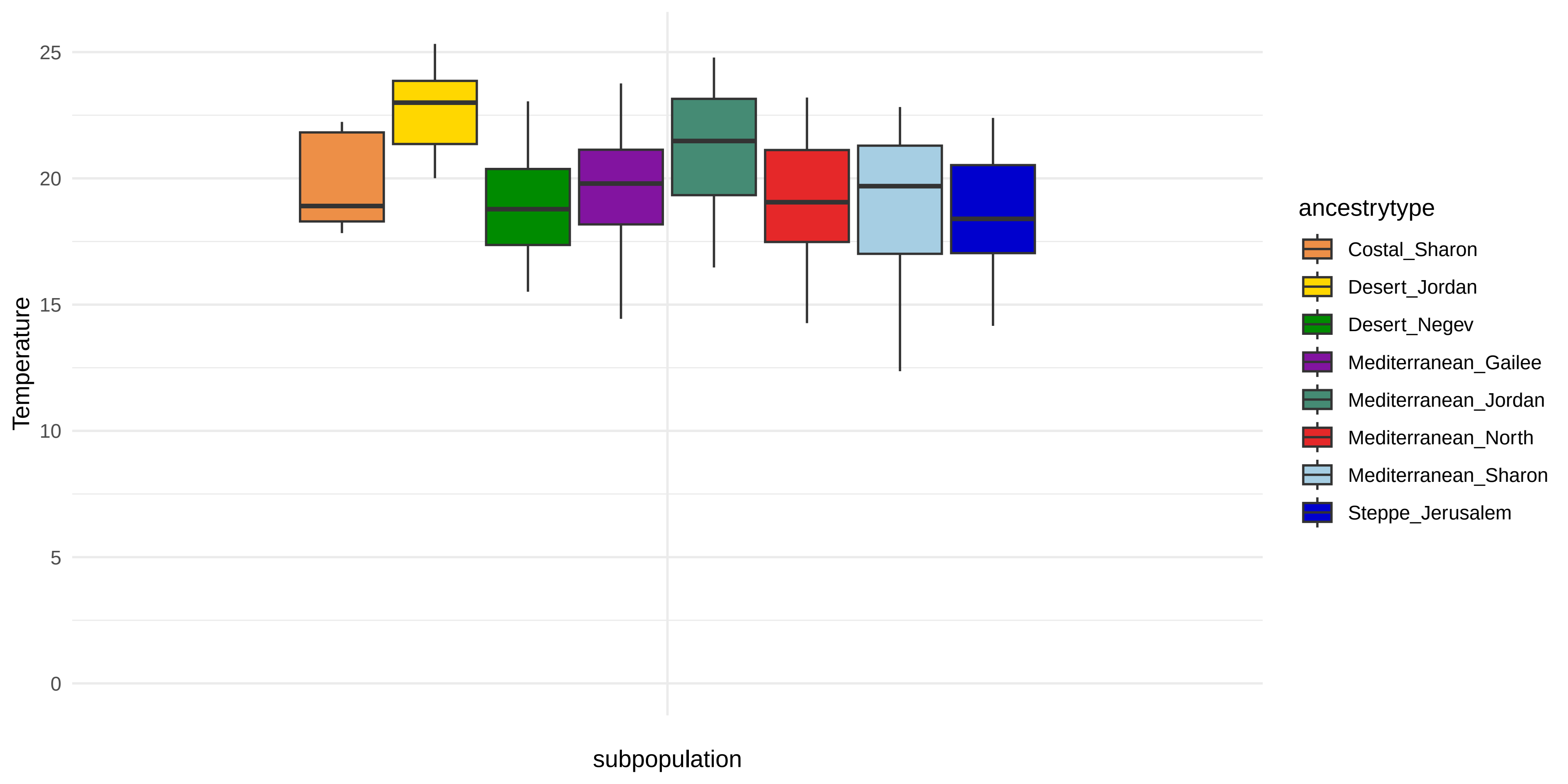

### Supplemental Figure S7

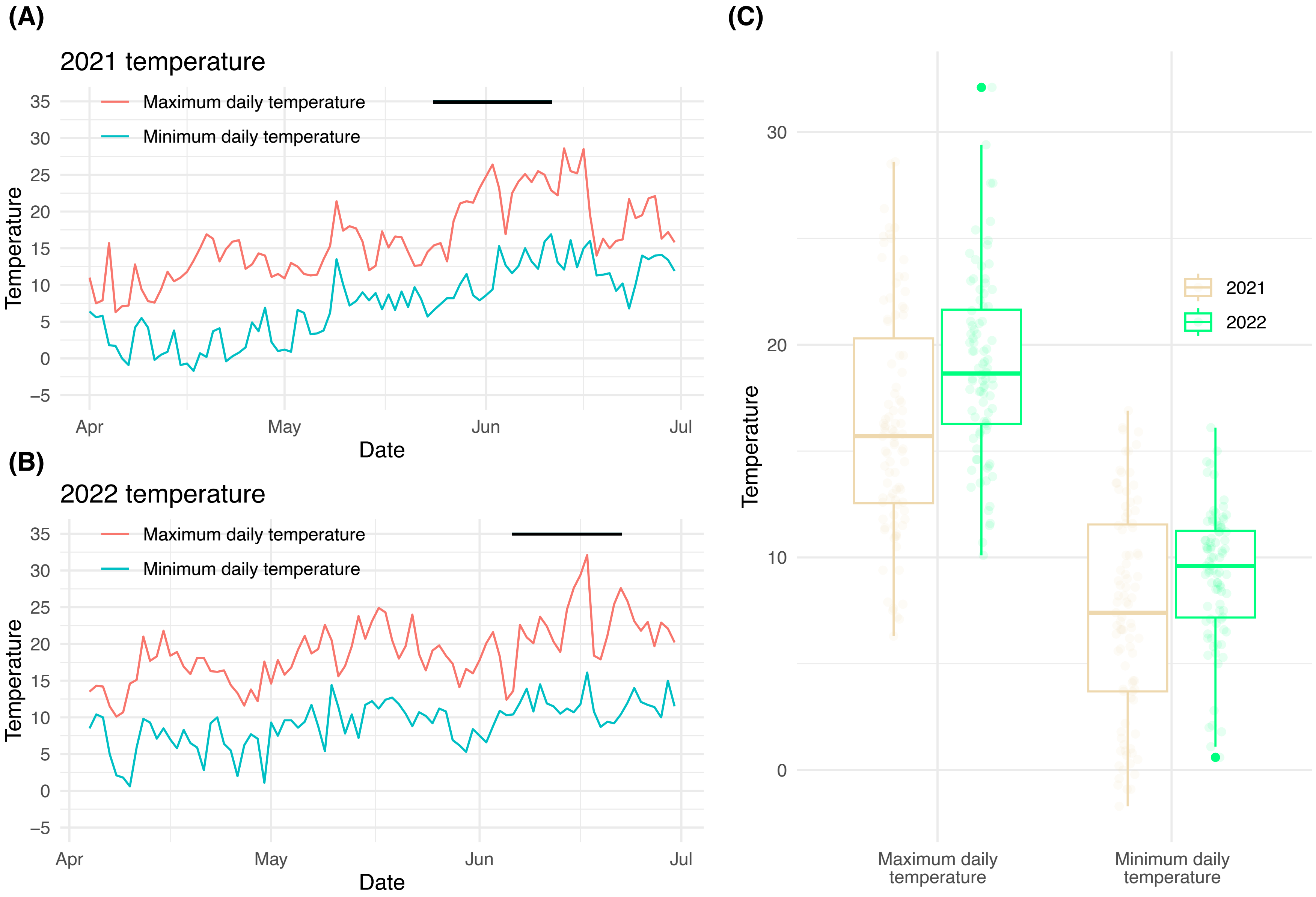

### Supplemental Figure S8

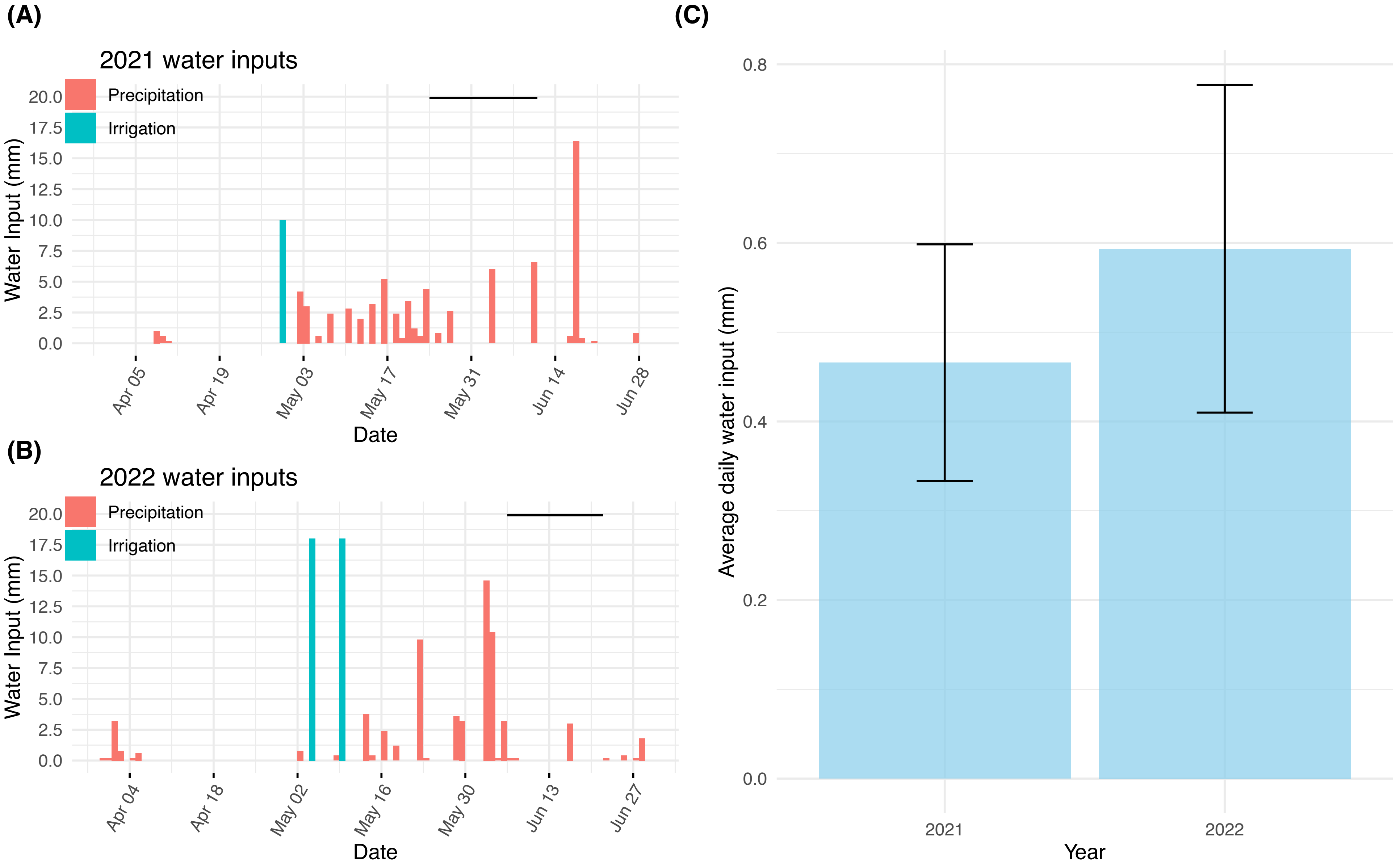

### Supplemental Figure S9

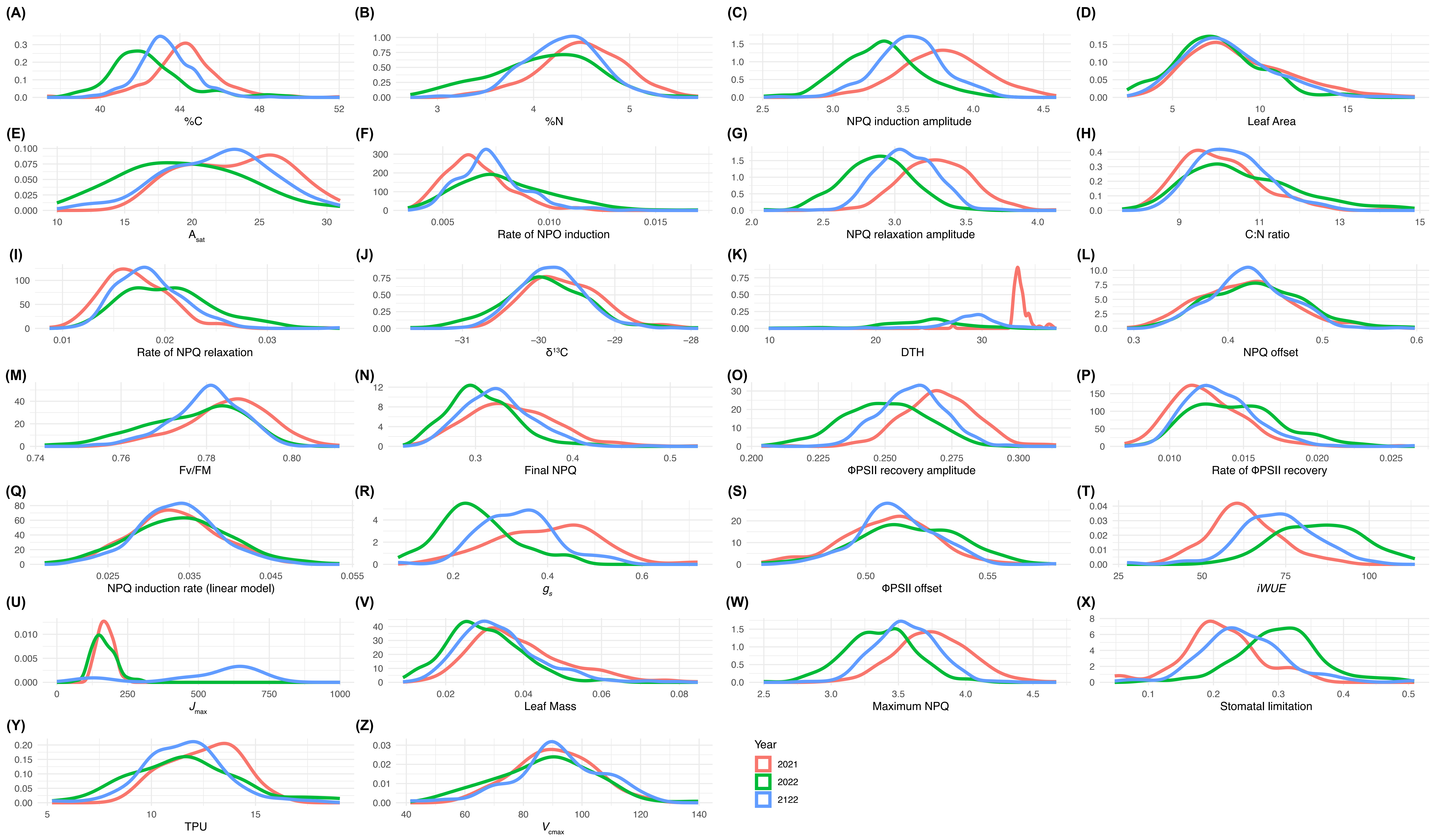

### Supplemental Figure S10

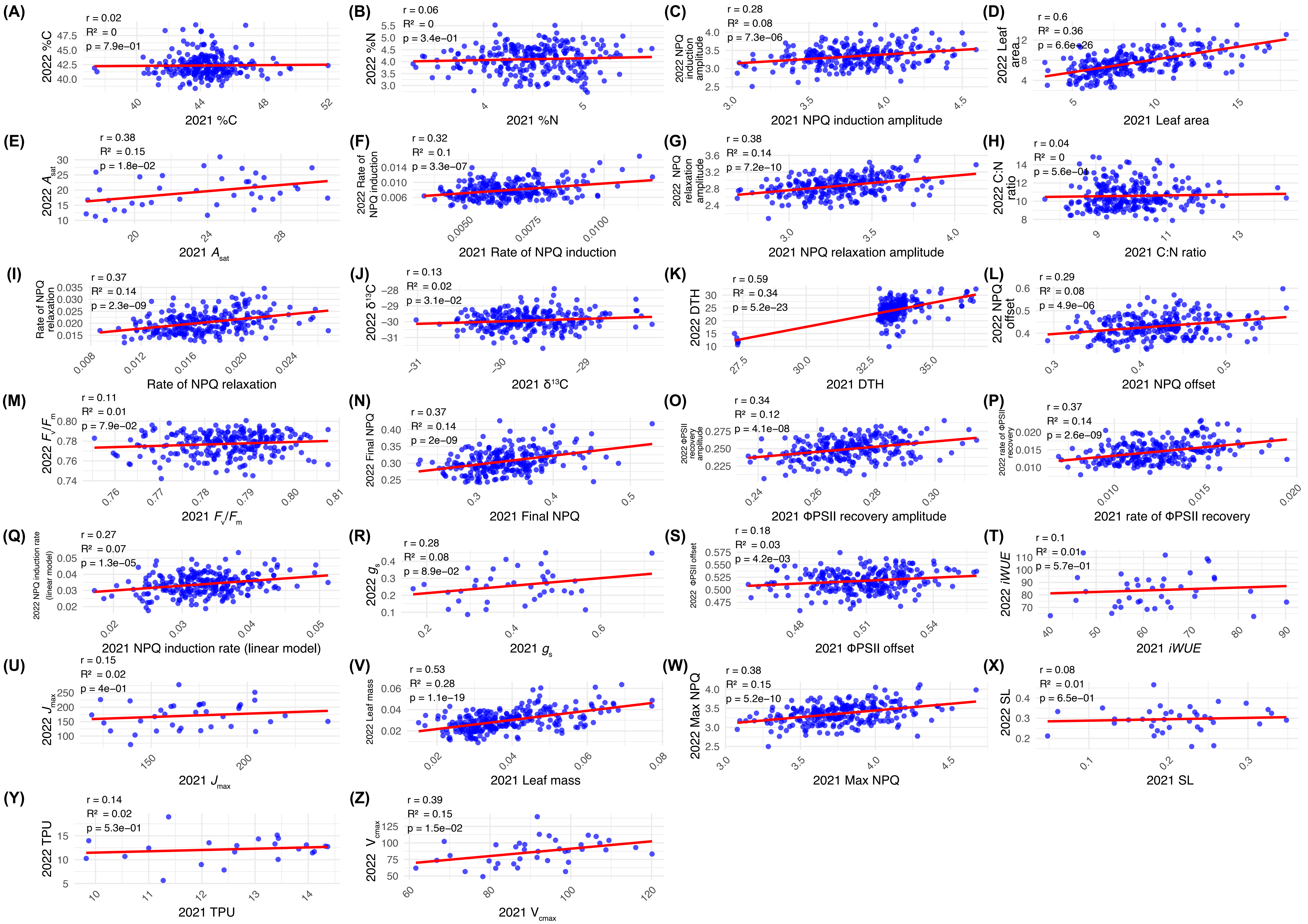

### Supplemental Figure S11

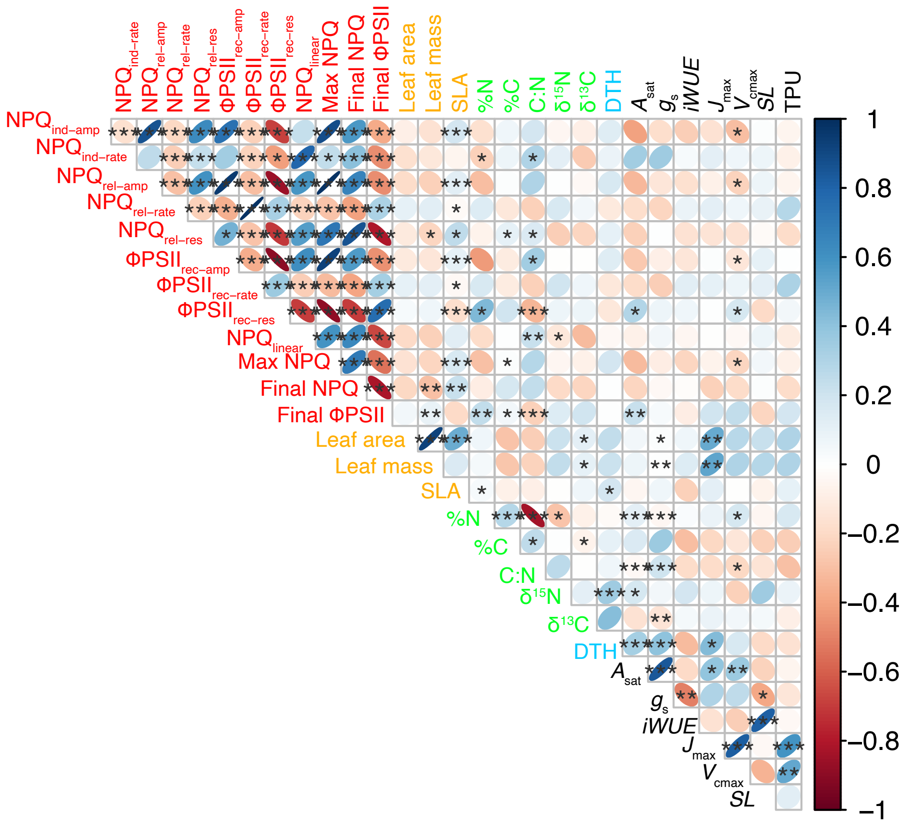

### Supplemental Figure S12

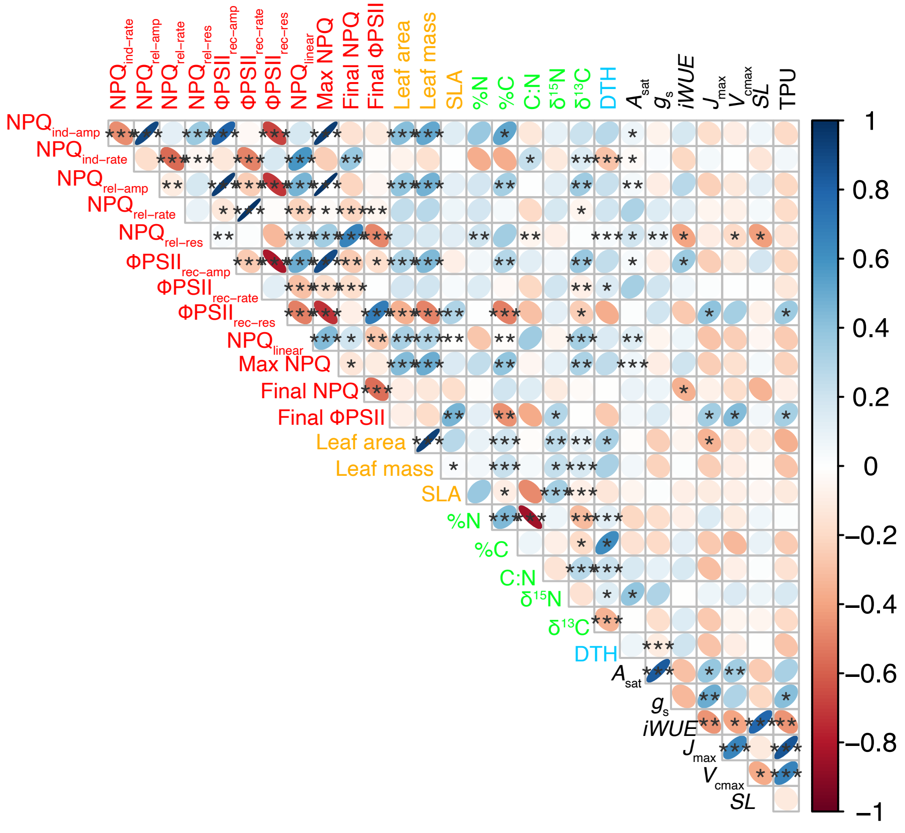

### Supplemental Figure S13

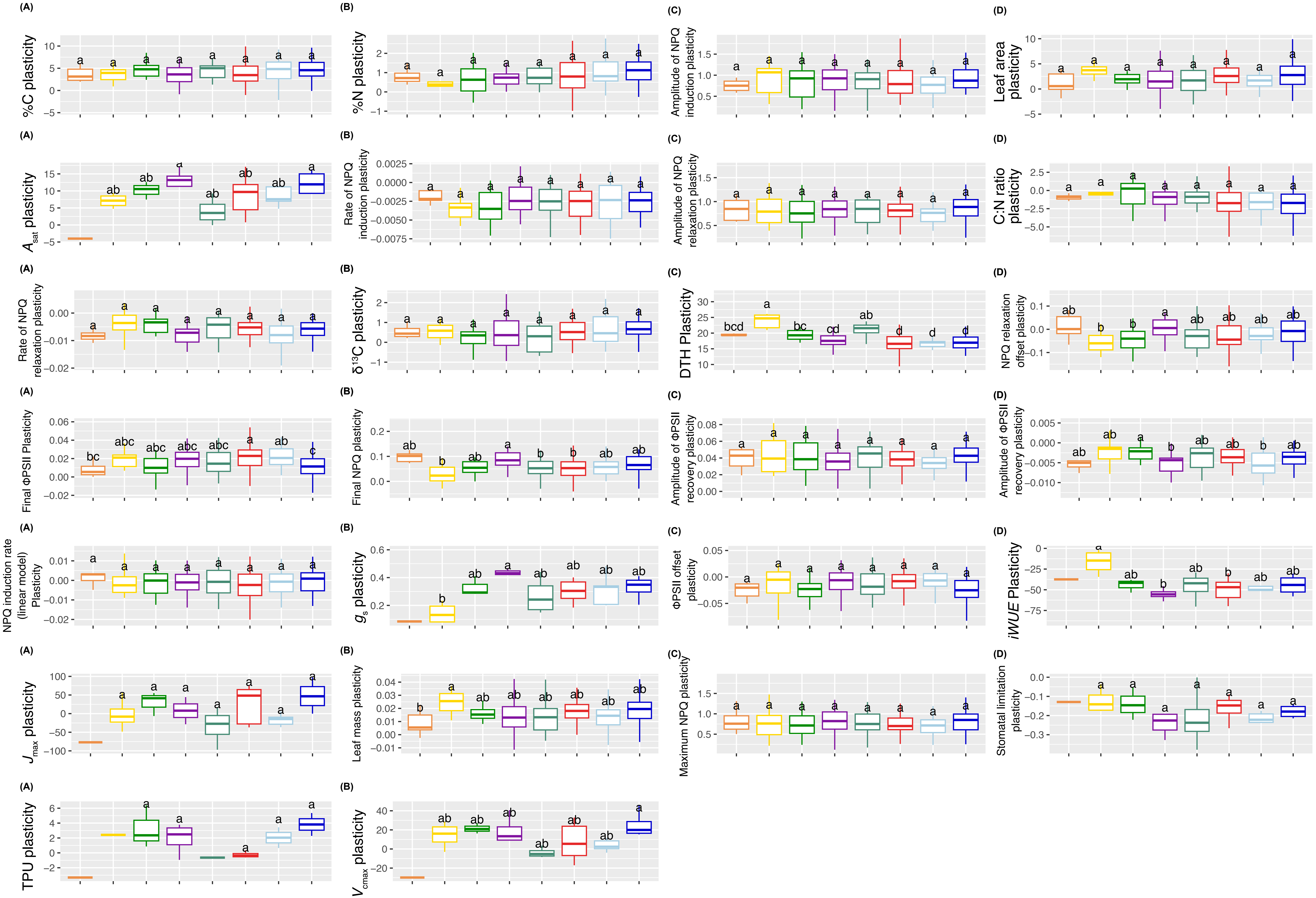

### Supplemental Figure S14

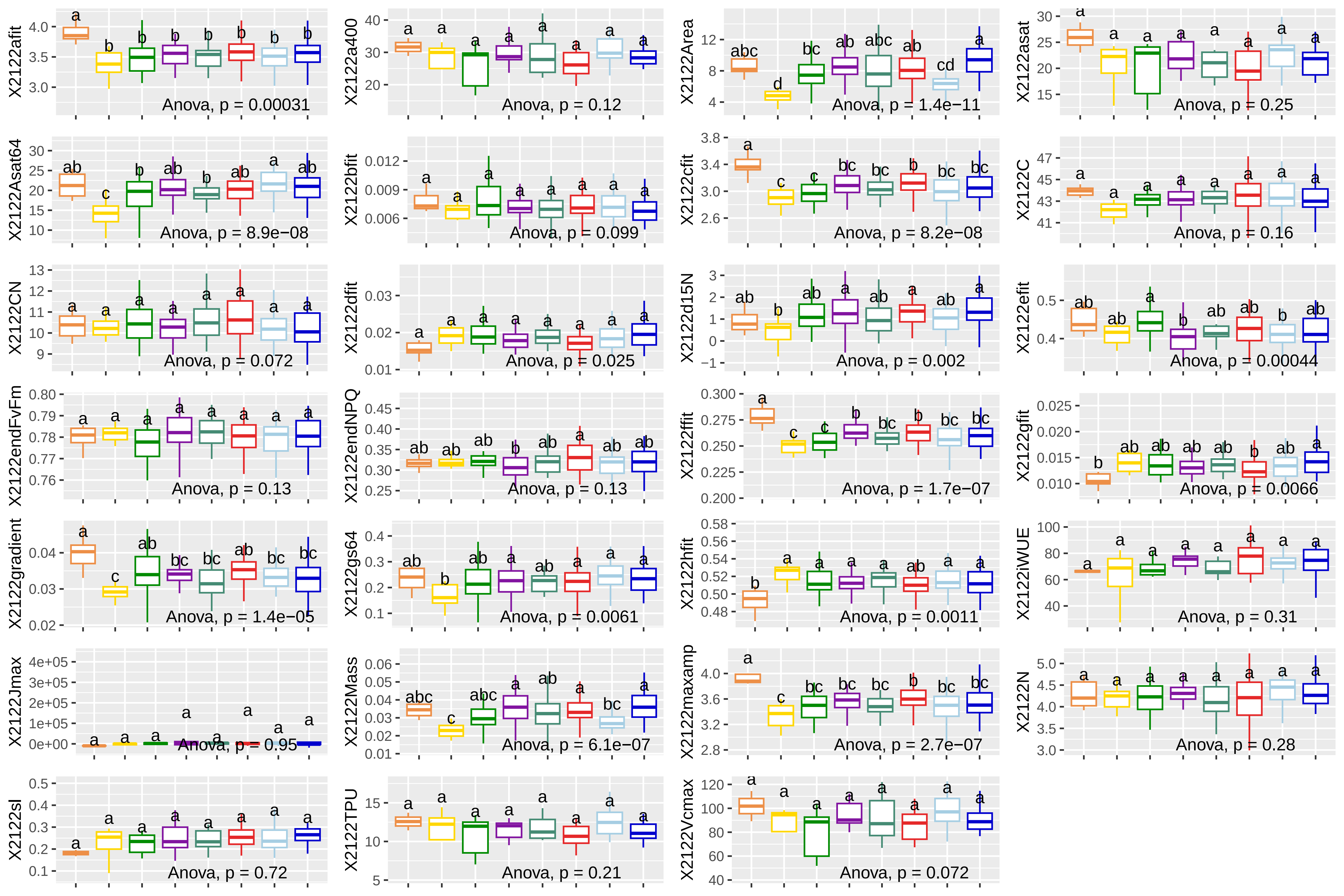
